# TBK1 and IKKε protect target cells from IFNγ-mediated T cell killing via an inflammatory apoptotic mechanism

**DOI:** 10.1101/2024.08.06.606693

**Authors:** Nicholas D. Sun, Allison R. Carr, Erica N. Krogman, Yogesh Chawla, Jun Zhong, Matthew C. Guttormson, Mark Chan, Michelle A. Hsu, Haidong Dong, Dusan Bogunovic, Akhilesh Pandey, Laura M. Rogers, Adrian T. Ting

## Abstract

Cytotoxic T cells produce interferon gamma (IFNγ), which plays a critical role in anti-microbial and anti-tumor responses. However, it is not clear whether T cell-derived IFNγ directly kills infected and tumor target cells, and how this may be regulated. Here, we report that target cell expression of the kinases TBK1 and IKKε regulate IFNγ cytotoxicity by suppressing the ability of T cell-derived IFNγ to kill target cells. In tumor targets lacking TBK1 and IKKε, IFNγ induces expression of TNFR1 and the Z-nucleic acid sensor, ZBP1, to trigger RIPK1-dependent apoptosis, largely in a target cell-autonomous manner. Unexpectedly, IFNγ, which is not known to signal to NFκB, induces hyperactivation of NFκB in TBK1 and IKKε double-deficient cells. TBK1 and IKKε suppress IKKα/β activity and in their absence, IFNγ induces elevated NFκB-dependent expression of inflammatory chemokines and cytokines. Apoptosis is thought to be non-inflammatory, but our observations demonstrate that IFNγ can induce an inflammatory form of apoptosis, and this is suppressed by TBK1 and IKKε. The two kinases provide a critical connection between innate and adaptive immunological responses by regulating three key responses: (1) phosphorylation of IRF3/7 to induce type I IFN; (2) inhibition of RIPK1-dependent death; and (3) inhibition of NFκB-dependent inflammation. We propose that these kinases evolved these functions such that their inhibition by pathogens attempting to block type I IFN expression would enable IFNγ to trigger apoptosis accompanied by an alternative inflammatory response. Our findings show that loss of TBK1 and IKKε in target cells sensitizes them to inflammatory apoptosis induced by T cell-derived IFNγ.

**Short Summary:** In the absence of TBK1 and IKKε, target cells are killed by T cells in an IFNγ-dependent manner. In TBK1 and IKKε-deficient cells, IFNγ induces RIPK1-dependent death, as well as hyper-induction of NFκB-dependent inflammatory genes. This suggests that any inhibition of TBK1/IKKε to block type I IFN expression will result in the demise of the cell accompanied by an alternate inflammatory program.

**Graphical Abstract:** 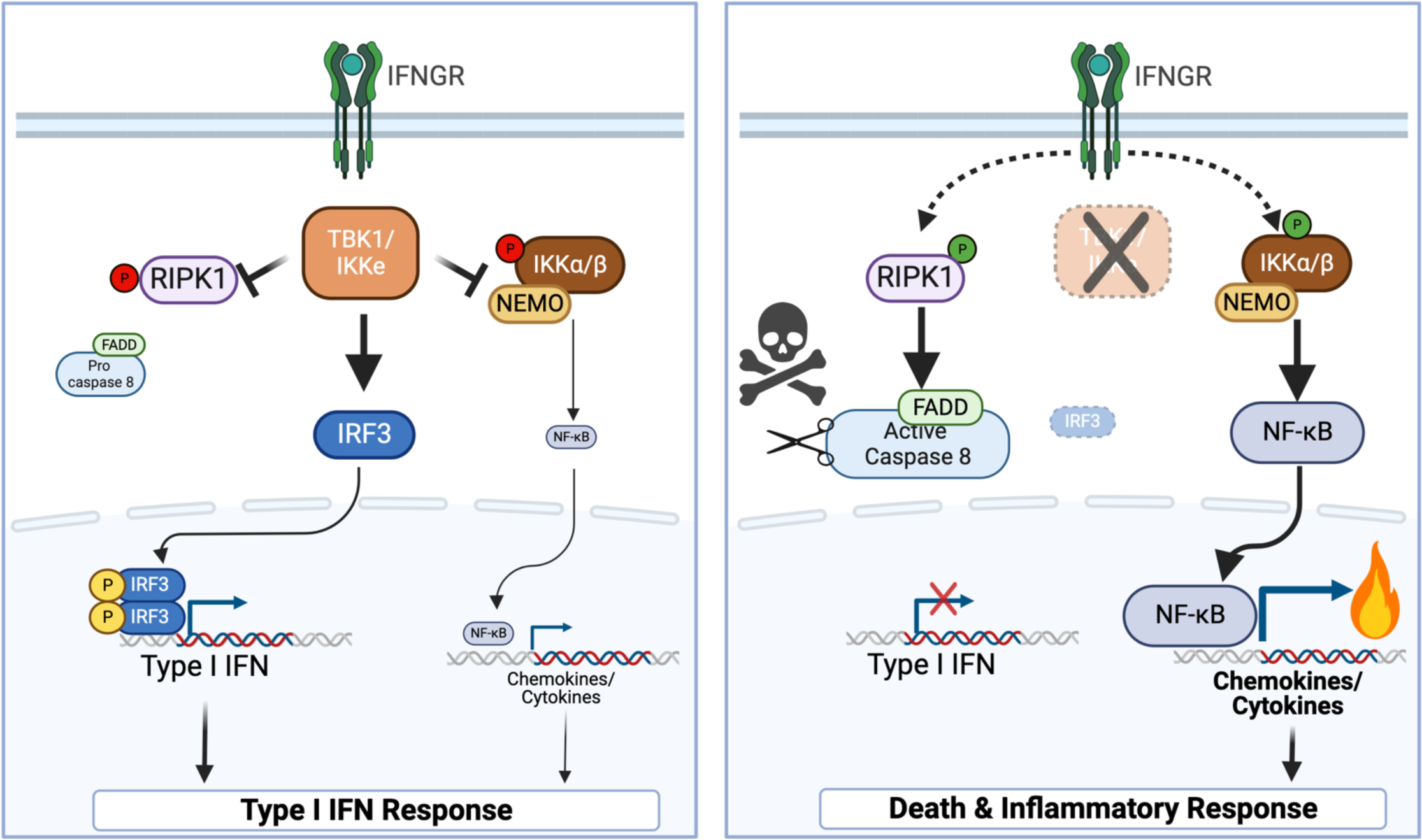

## Introduction

Cytotoxic lymphocytes, including CD8+ cytotoxic T lymphocytes (CTLs) and natural killer (NK) cells, are central to anti-viral and anti-tumor immunity. These cytotoxic cells kill target cells via their deposition of cytotoxic granules containing perforin and granzymes onto target cells, or engagement of FASL and TRAIL with their respective death receptors on the target cells. Activated CTLs and NK cells also produce cytokines including the hallmark Th1 cytokine, interferon gamma (IFNγ), which plays a critical role in anti-viral and anti-tumor immunity. This function of IFNγ has been shown to be due to its role in Th1 differentiation and activation of other immune cells. For instance, IFNγ induces the differentiation of CD8 CTLs^1, 2^ and enhances the antigen presentation machinery on antigen-presenting cells during the priming phase^3^ as well as on target cells during the effector phase^4^. However, it is not clear whether IFNγ can directly kill target cells engaged by T cells.

TANK-binding kinase 1 (TBK1) and I-kappaB kinase epsilon (IKKε) are two related kinases with redundant function^5, 6^ in multiple innate immune signaling pathways including TLRs, RLRs and cGAS. They phosphorylate the transcriptional factors IRF3 and IRF7, which is required to induce type I IFN expression following sensing of microbial infection by pattern recognition receptors^7, 8^. The two kinases were initially discovered during efforts to identify kinases that phosphorylate IκB molecules in the NFκB pathway. Deletion of *Tbk1* on the B6 background resulted in embryonic lethality whereas deletion of *Ikbke* (coding for IKKε) did not result in an overt phenotype^9, 10^. These genetic studies, including with cells deficient for both kinases, also showed that they were not essential for the activation of NFκB^6, 9^. Due to a prior study showing that defective NFκB activity (due to a deletion in p65 subunit *Rela*) enhanced sensitivity to TNF-induced cell death and resulted in embryonic lethality^11^, the lethal phenotype of *Tbk1^-/-^* mice with an intact NFκB signaling pathway was a surprising observation. In addition, since the embryonic lethality of *Tbk1^-/-^* mice could be rescued by a compound deletion of *Tnfrsf1a* (coding for TNFR1) or *Tnf*^5, 9^, this suggested that TBK1 phosphorylates a substrate in the TNFR1 pathway to block lethality. This substrate was subsequently shown to be RIPK1, whose phosphorylation by TBK1 and IKKε suppresses RIPK1’s death-signaling function in response to TNFR1 ligation^12, 13, 14, 15, 16^. Since the kinase activity of RIPK1 is required for its death-signaling function, the embryonic lethality of *Tbk1^-/-^* mice can also be reversed by the kinase-inactive *Ripk1^D138N^* allele^16^. In humans, a deficiency in TBK1 leads to chronic and systemic autoinflammation driven by elevated cell death that can be ameliorated by a TNF antagonist^17^. The loss or inhibition of TBK1 in tumor cells has also been reported to enhance the cytotoxicity of TNF and IFNγ^18^, but the signaling crosstalk between the two cytokines was not elucidated. While TBK1 and IKKε are well studied in type I IFN and TNF responses, a role for these kinases downstream of IFNγ is unknown.

Here, we report that target cell expression of TBK1 and IKKε protects against the cytotoxic effect of IFNγ. Deletion of TBK1 and IKKε in target cells sensitizes them to IFNγ-induced apoptosis mediated by cell-autonomous activation of TNFR1 and ZBP1. Paradoxically, apoptotic death in TBK1 and IKKε-deficient targets is accompanied by hyperactivation of canonical and non-canonical NFκB, indicating that TBK1 and IKKε are inhibitors of NFκB signaling. The elevated NFκB leads to potent induction of inflammatory chemokines and cytokines. These observations indicate that TBK1 and IKKε suppress IFNγ-induced inflammatory apoptosis.

## Results

### TBK1/IKKε-deficient cells succumb to IFNγ-mediated killing

We recently published that target cells that are deficient in the molecule SHARPIN are more susceptible to killing by T cells secreting TNF^19^. This causes SHARPIN-deficient organ transplants to be more easily rejected by allo-reactive T cells and SHARPIN-deficient B16-F1 (B16) melanoma cells to become more susceptible to immune checkpoint blockade with anti-PD1^19^. SHARPIN is a component of the linear ubiquitin assembly complex (LUBAC) E3 ligase that ubiquitinates RIPK1, and since TBK1/IKKε functions downstream of LUBAC^12^, we sought to study whether the two kinases played a similar role in regulating target cell sensitivity to TNF-dependent T cell killing. Using Crispr-Cas, we generated TBK1 and IKKε single knockouts (KO) in B16 cells, as well as a double knockout (DKO) of the two kinases due to their known redundancy ^5, 6^. Western blot analysis confirmed lack of protein expression of each kinase (Fig. 1A). Absence of phosphorylated IRF-3 upon poly(I:C) transfection confirmed the functional deficiency of TBK1 and IKKε in the DKO cells (Extended Data Fig. 1A). To test the sensitivity of these KO B16 lines to cytokine-induced cell death, we utilized the IncuCyte real time imaging system. Tumor killing events were quantified as a measure of YOYO-3 fluorescence counts normalized to the confluency taken at each time point. As expected, TBK1 KO and DKO cells were largely sensitive to TNF-induced killing, whereas control B16 cells transduced with a non-targeting sgRNA, hereafter referred to as wildtype (WT), displayed minimal cell death (Fig. 1B). IKKε KO showed significantly less death than TBK1 KO indicating a lesser role of the kinase, but lack of both kinases provided the greatest sensitivity to TNF. In our initial experiment, we also stimulated these cell lines with IFNγ as a negative control, as IFNγ signaling is coupled to the JAK-STAT pathway and not directly to any caspase-dependent pathway. Contrary to our expectations, while WT cells were fully resistant to IFNγ, both TBK1 KO and DKO exhibited significant sensitivity to IFNγ-mediated cell death, with DKO being the most sensitive (Fig. 1B). Of note, IFNγ stimulation induced similar level of STAT1 phosphorylation in the DKO cells, confirming the absence of the two kinases did not affect the proximal interferon gamma receptor (IFNGR) signaling pathway (Extended Data Fig. 1B). As deficiencies in other signaling molecules in the TNFR1 pathway are known to confer sensitivity to cell death in response to TNF, we sought to see if these deficiencies similarly conferred sensitivity to IFNγ-induced death. While B16 cells that lacked either SHARPIN^19, 20, 21^, TRAF2^22^ (component of the cIAP1/2 K63-linked ubiquitin E3 ligase), or both were significantly killed by TNF, these mutant cells were resistant to IFNγ with only minimal cell death seen in the SHARPIN/TRAF2 DKO cells (Extended Data Fig. 1C). Similarly, NEMO-deficient^23^ B16 cells were highly sensitive to TNF but were fully resistant to IFNγ (Extended Data Fig. 1D). These observations suggested that a deficiency in TBK1/IKKε, but not in other components of the TNFR1 pathway, uniquely sensitized cells to IFNγ-mediated death. Furthermore, type I IFN did not induce the death of DKO cells, suggesting that this is a type II IFN-specific response (Extended Data Fig. 1E).

**Figure 1.**
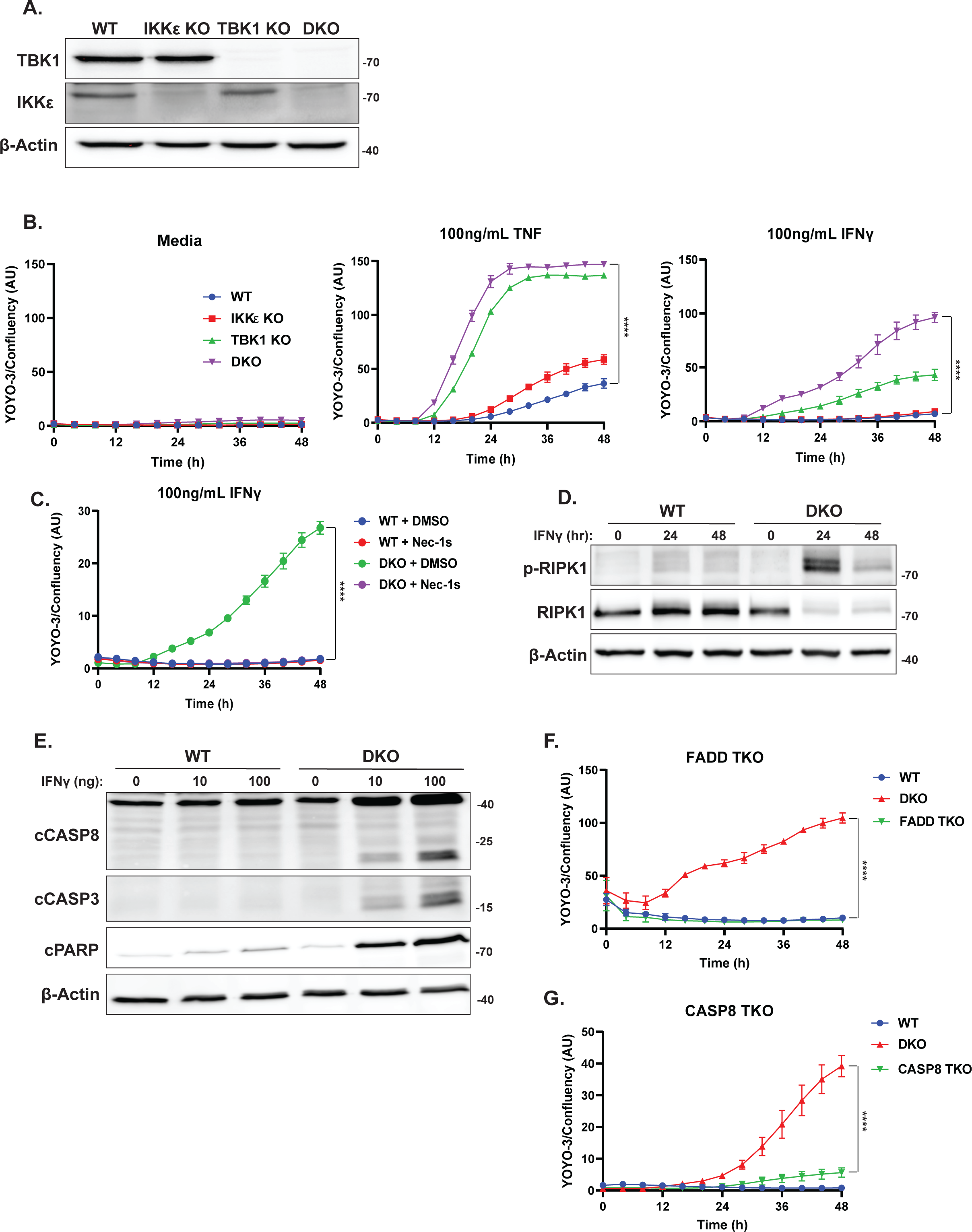
TBK1/IKKε-deficient cells are sensitive to IFNγ-induced RIPK1-mediated apoptosis. **(A)** Western blot analysis of the indicated proteins in wildtype (WT), IKKε single knockout, TBK1 single knockout, and double knockout (DKO) B16-F1 melanoma cells generated by CRISPR. **(B)** Control sgRNA (WT), IKKε and TBK1 single KO, and DKO B16 cell lines were stimulated in triplicates with media, 100 ng/mL TNF, or 100 ng/mL IFNγ and imaged for 48 h in an IncuCyte S3 in the presence of YOYO-3 to measure cell death. The confluency of the cells in each well were also quantified. Data is presented as YOYO-3 counts normalized to confluency in each well. Values are triplicate mean ± SD. ****P<0.0001 by unpaired t test comparing last data point. **(C)** WT and DKO B16 cells were treated with IFNγ (100 ng/mL) in the presence of DMSO or Nec-1s (10 μM) and analyzed in the IncuCyte. Values are triplicate mean ± SD. ****P<0.0001 by unpaired t test comparing last data point. **(D)** Western blot analysis of the indicated proteins in WT and DKO B16 cells pretreated with zVAD-fmk (20 μM) for 30 min, followed by IFNγ (100 ng/mL) for 0, 24, and 48 h. **(E)** Western blot analysis of the indicated proteins in WT and DKO B16 cells treated with 0, 10, 100 ng/mL IFNγ for 24 h. **(F-G)** WT, DKO, TBK1/IKKε/FADD TKO **(F)**, and TBK1/IKKε/CASP8 TKO **(G)** B16 cells were treated with IFNγ (100 ng/mL) and analyzed by IncuCyte. Values are triplicate mean ± SD. ****P<0.0001 by unpaired t test comparing last data point.

To further elucidate the mechanism underlying IFNγ-driven cell death in cells that lacked TBK1 and IKKε, we first assessed the importance of their kinase function by reconstituting the DKO cells with TBK1 WT, IKKε WT, or kinase-inactive TBK1 (TBK1 K38A). While the WT form of both kinases reversed the cell death sensitivity seen in the DKO cells, TBK1 K38A did not (Extended Data Fig. 1F, G). This observation suggests that TBK1 and IKKε can both protect against death, consistent with their known redundancy^5, 6^, and this function is dependent on the kinase activity. Considering their redundancy, and as the DKO exhibited the strongest phenotype, our subsequent analysis was carried out using the DKO cells. As RIPK1 is known to be required for TNF-induced cell death in TBK1-deficient cells^12, 16^, we next examined to see if it was similarly involved in the IFNγ response. Necrostatin-1s (Nec-1s), an inhibitor of the kinase activity of RIPK1, inhibited the cell death in the DKO cells induced by IFNγ (Fig. 1C). In addition, we examined the phosphorylation of RIPK1 on Ser166, a marker of RIPK1 death-signaling^24^, in WT, DKO, and the reconstituted cells after IFNγ stimulation. We observed markedly enhanced phosphorylation of RIPK1 on Ser166 in the DKO and TBK1 K38A cells (Fig. 1D and Extended Data Fig. 1H). RIPK1 Ser166 phosphorylation was accompanied by a decrease of RIPK1 protein in the detergent-soluble compartment (Fig. 1D), which is another biochemical hallmark of RIPK1-dependent death^25, 26^. Collectively, these data suggest the death of the DKO B16 cells induced by IFNγ is dependent on RIPK1. To determine which form of cell death IFNγ is triggering in the DKO cells, we performed western blotting with antibodies against cleaved CASP8, CASP3, and PARP. These biochemical hallmarks of apoptosis were induced by IFNγ in the DKO cells, but not when the DKO cells were complemented with TBK1 WT (Fig. 1E and Extended Data Fig. 1I). IFNγ-induced death of DKO cells was dependent on FADD and CASP8 as it was abrogated in TBK1/IKKε/FADD-deficient (FADD TKO) and TBK1/IKKε/CASP8-deficient (CASP8 TKO) cells, respectively (Fig. 1F, G). The observation that cell death of the DKO cells was completely reversed by the absence of FADD suggested that necroptosis is not induced in the DKO cells. This suggestion was further supported by the observation that B16 cells do not express RIPK3, which is essential for necroptosis^27, 28, 29^ (Extended Data Fig. 1J). We further examined the phosphorylation of MLKL on Ser345, a biochemical hallmark of necroptosis^30^. IFNγ did not induce detectable MLKL phosphorylation whereas this was observed in the positive control of mouse embryonic fibroblasts (MEF) stimulated with a combination of TNF, SMAC mimetic and zVAD-fmk (Extended Data Fig. 1K). While we cannot rule out necroptosis occurring at a level below our detection limit, our observations indicate that apoptosis is the dominant form of cell death and led us to conclude that IFNγ induces RIPK1-dependent apoptosis in TBK1/IKKε-deficient tumor cells.

### TBK1/IKKε-deficient target cells are killed by T cells in an IFNγ-dependent manner

Since effector CD8 T cells is a major producer of IFNγ, we next asked whether T cells can utilize this cytokine to kill TBK1/IKKε-deficient targets. To assess this, we pulsed WT or DKO B16 cells with control GP33 or OVA SIINFEKL peptide and co-cultured them with OT-I CD8 T cells for 48 hours. We found that even at a low T cell effector to target (E/T) ratio, the DKO targets were highly sensitive to OT-I killing only when the OVA antigen was present, while the WT targets, regardless of antigen, were resistant to killing (Fig. 2A). Furthermore, the reconstitution of TBK1 WT into the DKO cells protected them from OT-I T cell-mediated killing, while TBK1 K38A failed to do so (Extended Data Fig. 2A). Similar to recombinant IFNγ stimulation, OT-I killing of DKO targets is dependent on FADD, CASP8 and RIPK1 kinase activity in the target cells (Fig. 2B, C and Extended Data Fig. 2B). To confirm whether IFNγ, or potentially TNF, secreted by T cells is killing the DKO targets, we added blocking antibodies against IFNγ or TNF to the OT-I and DKO co-cultures. Interestingly, only anti-IFNγ was able to block the cell death mediated by the T cells, whereas anti-TNF was largely ineffective (Fig. 2D). ELISA of supernatants from the OT-I and tumor target co-cultures confirmed that the T cells are producing both cytokines, though IFNγ levels appear to be higher than TNF in the supernatants (Extended Data Fig. 2C). Since CD8 T cells can also utilize perforin and granzyme to kill target cells^31^, we wanted to rule this role out as a potential cytotoxic mechanism in the killing of the DKO targets. We transduced the OT-I TCR α and β chains^32^ into WT or perforin KO (PRF1 KO) T cells and co-cultured them with the DKO targets. Both WT and PRF1 KO T cells killed OVA-pulsed DKO targets equally well (Extended Data Fig. 2D). These observations demonstrate that TBK1/IKKε-deficient target cells are sensitized to T cell killing mediated by IFNγ but not perforin. To determine if loss of TBK1/IKKε in target cells enhanced their killing by T cells *in vivo*, we first transfected the cytoplasmic ovalbumin (cOVA) gene into B16 WT and DKO cells (WT-cOVA and DKO-cOVA). When co-cultured with OT-I T cells, the DKO-cOVA targets, but not the WT-cOVA targets, were killed, confirming functional OVA expression (Extended Data Fig. 2E). When target cells were implanted into NOD.Cg-*Prkdc^scid^il2rg^tm1Wjl^*/SzJ (NSG) mice, tumor growth was comparable between mice with WT-cOVA and DKO-cOVA tumors when treated with PBS, whereas adoptive transfer of OT-I T cells resulted in better control of TBK1/IKKε-deficient tumors than WT control tumors (Fig. 2E). These results confirm that TBK1/IKKε-deficient target cells also have enhanced sensitivity to T cell-mediated killing *in vivo*. Finally, to see if these kinases also protect against IFNγ-induced apoptosis in a different cell line, we examined SVEC4-10 endothelial tumor cells. SVEC4-10 cells express TBK1 but no detectable level of IKKε compared to the positive control RAW264.7 macrophages (Extended Data Fig. 2F). Similar to B16 cells, deletion of *Tbk1* in SVEC4-10 cells was sufficient to confer sensitivity to IFNγ-induced apoptosis (Extended Data Fig. 2G, H).

**Figure 2.**
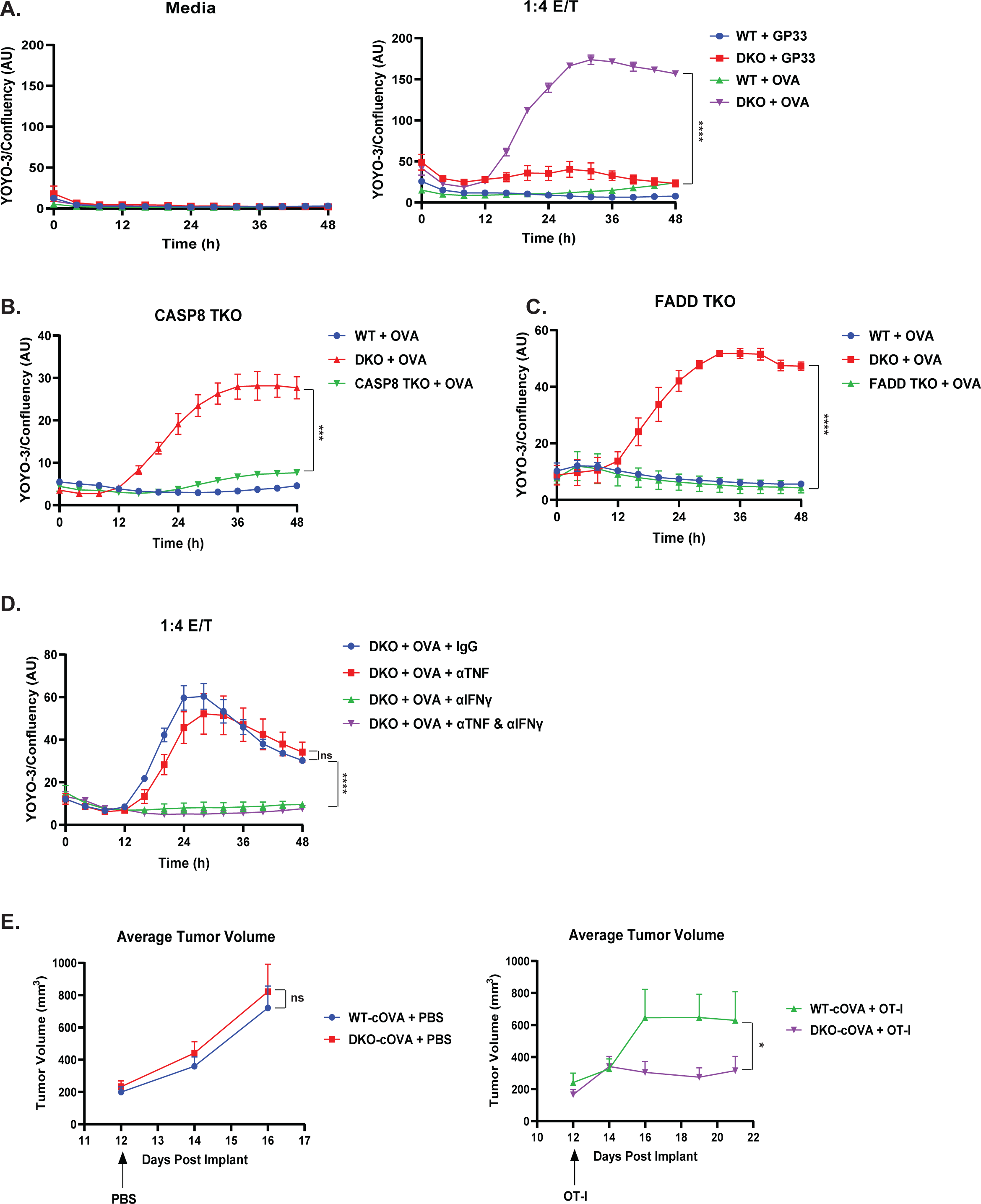
TBK1/IKKε-deficient targets are killed by T cells in an IFNγ-dependent manner. **(A)** WT and DKO B16 cells were pulsed with control LCMV GP33 peptide or OVA SIINFEKL peptide followed by co-culture with either control media or OT-I T cells at 1:4 effector to target (E/T) ratio. Target cell death was analyzed by IncuCyte. Values are triplicate mean ± SD. ****P<0.0001 by unpaired t test comparing last data point. **(B-C)** WT, DKO, TBK1/IKKε/CASP8 TKO **(B)**, and TBK1/IKKε/FADD TKO **(C)** B16 cells were pulsed with OVA peptide followed by co-culture with OT-I T cells at 1:4 E/T ratio. Target cell death was analyzed by IncuCyte. Values are triplicate mean ± SD. ***P<0.001, ****P<0.0001 by unpaired t test comparing last data point. **(D)** DKO B16 cells were pulsed with OVA peptide followed by co-culture with OT-I T cells at 1:4 E/T ratio and treated with control IgG (50 μg/mL), anti-TNF (50 μg/mL), anti-IFNγ (50 μg/mL) or both mAbs (50 μg/mL). Values are triplicate mean ± SD. ****P<0.0001, not significant (ns) by unpaired t test comparing last data point. **(E)** Tumor volume analysis of NSG mice bearing B16 WT- or DKO-cOVA tumors treated once at day 12 post tumor implant with PBS or OT-I T cells (10 million/100 μL) i.v.; WT (PBS) *n* = 13, DKO (PBS) *n* = 11, WT (OT-I) *n* = 7, DKO (OT-I) *n* = 8. Values are mean ± SEM. Not significant (ns), *P<0.05 by paired t test comparing all combined data points.

### IFNγ-mediated killing of TBK1/IKKε-deficient target cells is TNFR1-dependent

Since it is unclear what role TBK1 and IKKε may have downstream of IFNGR, we first sought to validate that IFNγ was signaling through STAT1 to kill the DKO cells. To this end, we additionally knocked out STAT1 on top of the DKO cells creating TBK1/IKKε/STAT1-deficient cells (STAT1 TKO) (Extended Data Fig. 3A). While STAT1 TKO cells remain sensitive to TNF-induced killing, they are now resistant to killing by both recombinant IFNγ and OT-I T cells (Fig. 3A and Extended Data Fig. 3B, C). Treatment with ruxolitinib, a JAK1 and JAK2 inhibitor, also rescued the DKO cells from IFNγ-mediated killing (Fig. 3B). These observations strongly suggest that STAT1-dependent transcription may be needed for IFNγ to kill TBK1/IKKε-deficient B16 cells.

**Figure 3.**
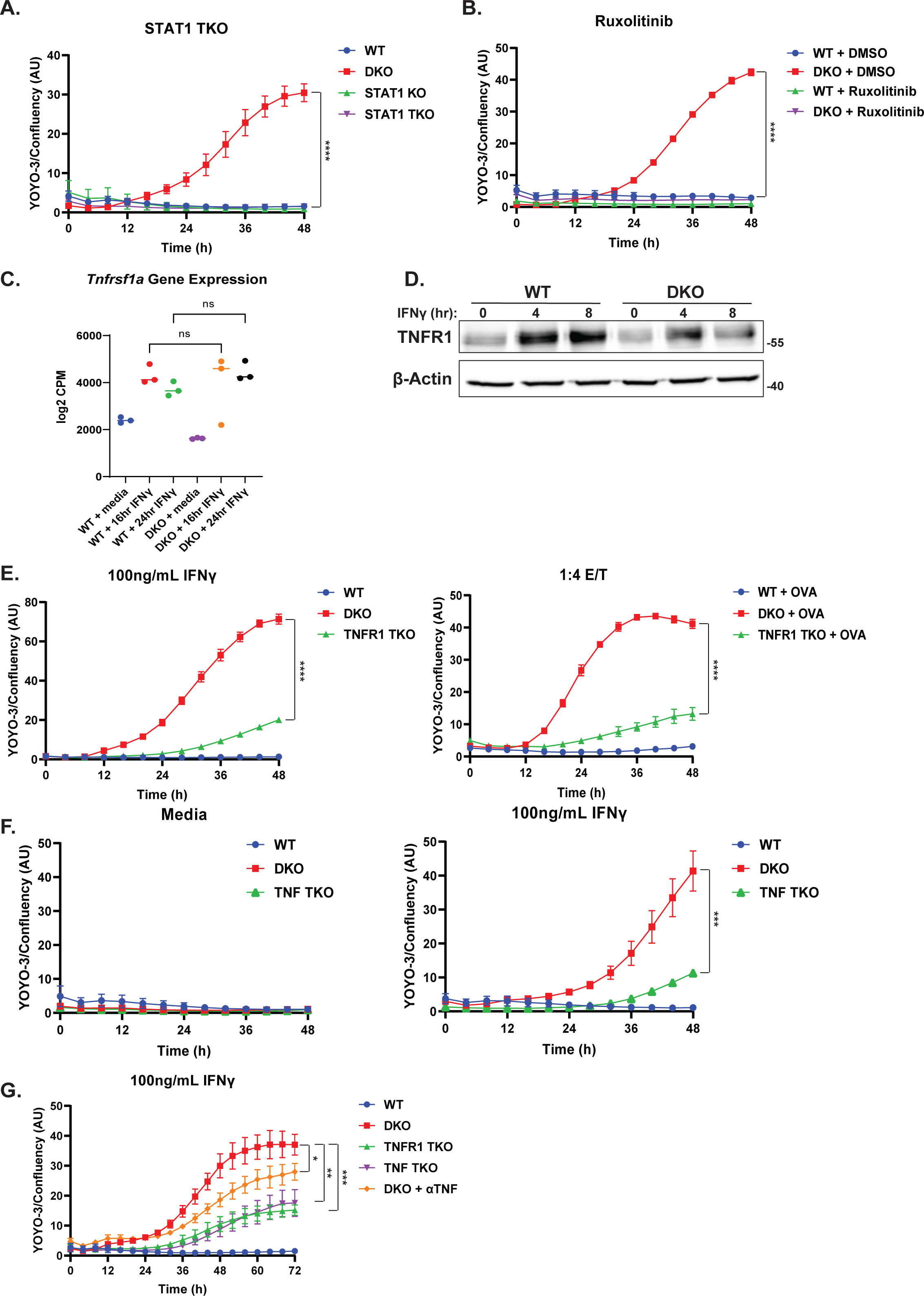
TNF/TNFR1 signaling is required for IFNγ-mediated killing of TBK1/IKKε-deficient cells. **(A)** WT, DKO, STAT1 single KO, and TBK1/IKKε/STAT1 TKO B16 cell were treated with IFNγ (100 ng/mL) and analyzed by IncuCyte. Values are triplicate mean ± SD. ****P<0.0001 by unpaired t test comparing last data point. **(B)** WT and DKO B16 cells were treated with IFNγ (100 ng/mL) in the presence of DMSO or ruxolitinib (1 μM) and analyzed by IncuCyte. Values are triplicate mean ± SD. ****P<0.0001 by unpaired t test comparing last data point. **(C)** WT and DKO B16 cells were stimulated with IFNγ (100 ng/mL) for 0, 16, 24 h and RNA isolated for sequencing. Values displayed as log2 CPM comparing *Tnfrsf1a* from three independent experiments. Statistical analysis was performed using one-way ANOVA with Sidak’s multiple-comparison test. ns = not significant. **(D)** Western blot analysis of the indicated proteins in WT and DKO B16 cells treated with IFNγ (100 ng/mL) for 0, 4, 8 h. **(E)** WT, DKO, and TBK1/IKKε/TNFR1 TKO B16 cells were treated with IFNγ (100 ng/mL) (left) or pulsed with OVA peptide followed by co-culture with OT-I T cells at 1:4 E/T ratio (right) and analyzed by IncuCyte. Values are triplicate mean ± SD. ****P<0.0001 by unpaired t test comparing last data point. **(F)** WT, DKO, and TBK1/IKKε/TNF TKO B16 cells were treated with media alone or IFNγ (100 ng/mL) and analyzed by IncuCyte. Values are triplicate mean ± SD. ***P<0.001 by unpaired t test comparing last data point. **(G)** WT, DKO, TBK1/IKKε/TNFR1 TKO, TBK1/IKKε/TNF TKO, and DKO with anti-TNF (50 μg/mL) B16 cells were treated with IFNγ (100 ng/mL) and analyzed by IncuCyte. Values are triplicate mean ± SD. *P<0.05, **P<0.01, ***P<0.001 by unpaired t test comparing last data point.

To uncover which genes and/or pathways are involved in the killing of DKO cells by IFNγ, we performed bulk RNA-seq analysis on WT and DKO B16 cells stimulated with IFNγ. Since the death we observed was RIPK1-dependent (Fig. 1C), we focused on candidate genes that are known to signal via TBK1/IKKε and were reported to couple to RIPK1. These analyses revealed that TNFR1 gene expression was induced by IFNγ stimulation in both WT and the DKO cells (Fig. 3C). Western blot analysis confirmed increased TNFR1 protein expression in both WT and DKO cells upon IFNγ treatment (Fig. 3D). Strikingly, knocking out TNFR1 on top of the DKO cells (TNFR1 TKO) significantly protected the cells from IFNγ and OT-I T cell-induced killing (Fig. 3E). Of note, TNFR1 TKO was no longer sensitive to TNF treatment, confirming a functional deficiency in TNFR1 (Extended Data Fig. 3D, E). These results suggested the enhanced sensitivity to IFNγ in B16 cells lacking TBK1/IKKε is TNFR1-dependent and therefore may be dependent on autocrine TNF. This possibility was supported by the RNA-seq analysis, which showed that TNF gene expression was induced by IFNγ in the DKO cells (Extended Data Fig. 3F). However, we detected minimal soluble TNF in the culture supernatants from both IFNγ stimulated WT and DKO cells by ELISA (Extended Data Fig. 3G), suggesting the level of TNF may be low and/or it remained membrane-bound. We then knocked out TNF in the DKO cells (TNF TKO) (Extended Data Fig. 3H), which reversed the sensitivity of the DKO to cell death induced by IFNγ to the same degree as TNFR1 TKO (Fig. 3F, G). Interestingly, while a soluble antagonist antibody against TNF was also able to reduce IFNγ-induced death of DKO, it had a smaller effect than either the genetic knockout of TNFR1 or TNF (Fig. 3G). This suggests that TNF can activate TNFR1 signaling in a cis-manner. This cell-autonomous behavior of TNF has been previously reported in myeloid cells^33, 34^. Altogether, these data strongly suggest that IFNγ induces the upregulation and activation of the TNF/TNFR1 signaling axis to induce apoptosis in TBK1/IKKε-deficient cells in a cell-autonomous manner.

### IFNγ-induced ZBP1 acts in tandem with TNFR1 to kill TBK1/IKKε-deficient targets

Although we established TNFR1 as a key driver of IFNγ-mediated killing of TBK1/IKKε-deficient cells, we noticed both TNF and TNFR1 TKO cells were not completely protected from IFNγ stimulation (Fig. 3E, F). This led us to investigate what additional receptors might be contributing to the IFNγ sensitivity of the DKO cells. It has been reported that TNFR1 and Z-DNA binding protein 1 (ZBP1, also known as DAI or DLM-1) can act together to cause intestinal inflammation and necroptotic cell death when FADD and CASP8 signaling is impaired^35^. ZBP1 is a well-characterized cytoplasmic sensor of Z-nucleic acids. It contains a RIP homotypic interaction motif (RHIM) domain that interacts with RIPK3 and RIPK1 to mediate necroptosis and apoptosis in response to both exogenous and endogenous Z-RNA^36, 37, 38^. Furthermore, the expression and activity of ZBP1 is known to be regulated by IFNγ^39^. We therefore postulated that ZBP1 may be working in tandem with TNFR1 to trigger IFNγ-induced cell death in TBK1/IKKε-deficient cells. To assess the role of ZBP1, we examined if ZBP1 can interact with RIPK1 in our B16 cells. Co-immunoprecipitation showed that ZBP1 associates with RIPK1 upon IFNγ stimulation in both WT and DKO cells (Fig. 4A), though this signal was reduced in the DKO cells, likely due to the known translocation of the RIPK1 signaling complex to a detergent-insoluble compartment when RIPK1 is activating cell death. We then generated TBK1/IKKε/ZBP1-deficient (ZBP1 TKO) and TBK1/IKKε/TNFR1/ZBP1-deficient (QKO) cells to examine the relative contribution of TNFR1 and ZBP1 to IFNγ-mediated apoptosis of the DKO cells (Extended Data Fig. 4A). ZBP1 single deficiency provided no protection in the DKO cells as compared to the greater protection provided by the TNFR1 single deficiency (Fig. 4B). Most importantly, QKO cells were completely resistant to IFNγ-mediated killing (Fig. 4B). Since ZBP1 senses endogenous Z-RNA^40^, we also tested the effect of reducing the availability of this ligand by overexpressing the p150 isoform of ADAR1 in the TBK1/IKKε/TNFR1 TKO cells, which can disrupt Z-RNA base-pairing^41, 42^ (Extended Data Fig. 4B). This had the same effect as ZBP1 deletion (Extended Data Fig. 4C). In line with the IncuCyte data, induction of Ser166 phosphorylation on RIPK1 and apoptotic markers in the DKO cells were more inhibited by TNFR1 deficiency than by ZBP1 deficiency, and completely inhibited by TNFR1/ZBP1 double deficiency (Fig. 4C, D). Lastly, when these same cells were co-cultured with OT-I T cells, the ZBP1 TKO cells were not protected whereas the QKO cells were completely protected from T cell-mediated killing (Fig. 4E). Thus, in the absence of TBK1 and IKKε, target cells are killed by T cells producing IFNγ, which activates both TNFR1 and ZBP1 to induce RIPK1-dependent apoptosis.

**Figure 4.**
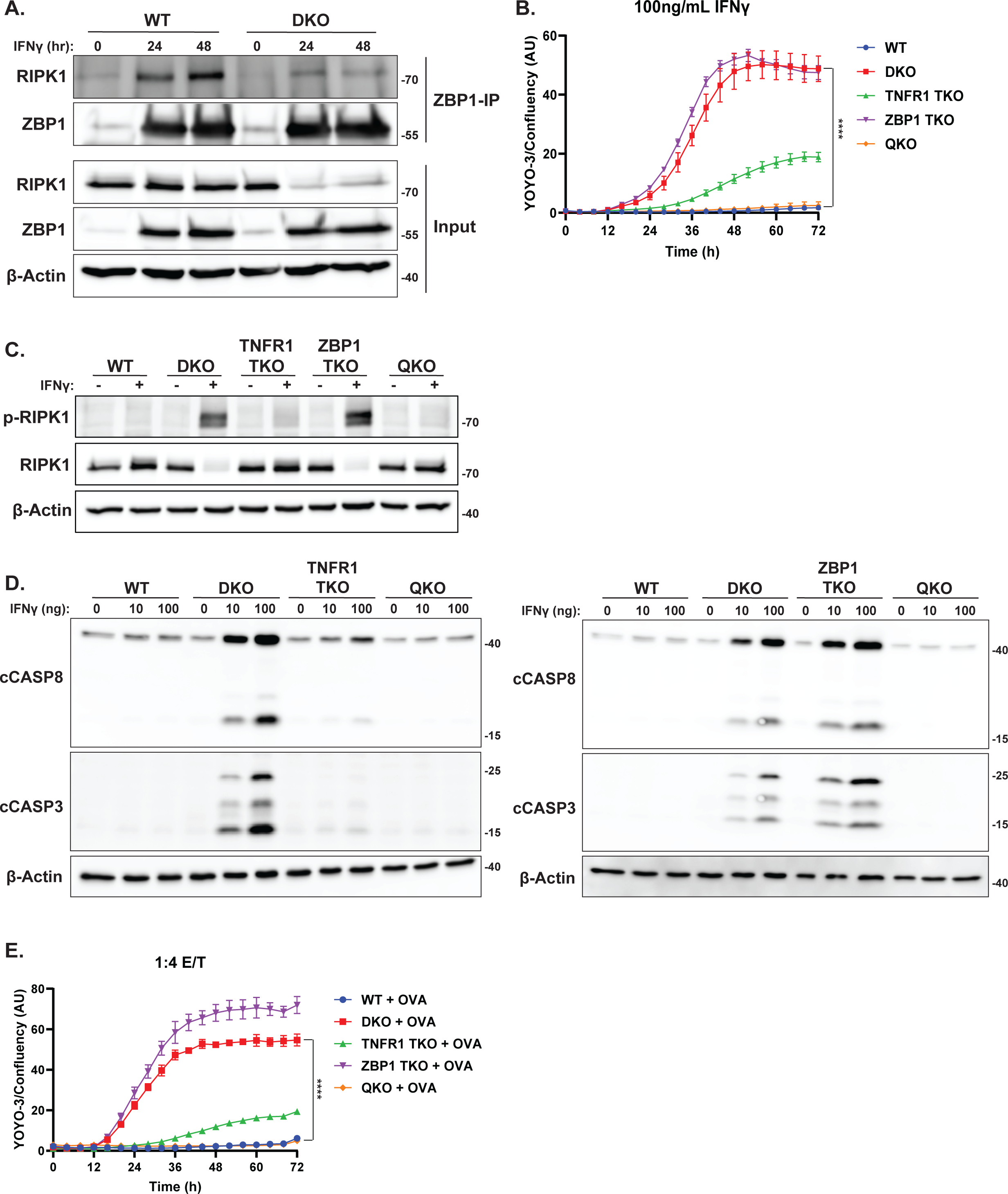
ZBP1 in tandem with TNFR1 mediate IFNγ killing of TBK1/IKKε-deficient cells. **(A)** Western blot analysis of the indicated proteins in WT and DKO B16 cells pretreated with zVAD-fmk (20 μM) for 30 min, followed by IFNγ (100 ng/mL) for 0, 24, and 48 h. Equal concentration of lysates were then immunoprecipitated with antibody against ZBP1. **(B)** WT, DKO, TBK1/IKKε/TNFR1 TKO, TBK1/IKKε/ZBP1 TKO, and TBK1/IKKε/TNFR1/ZBP1 QKO B16 cells were treated with IFNγ (100 ng/mL) and analyzed by IncuCyte. Values are triplicate mean ± SD. ****P<0.0001 by unpaired t test comparing last data point. **(C)** Western blot analysis of the indicated proteins in WT, DKO, TNFR1 TKO, ZBP1 TKO, and QKO B16 cells pretreated with zVAD-fmk (20 μM) for 30 min, followed by IFNγ (100 ng/mL) for 0 and 24 h. **(D)** Western blot analysis of the indicated proteins in WT, DKO, TNFR1 TKO, ZBP1 TKO, and QKO B16 cells treated with 0, 10, or 100 ng/mL IFNγ for 24 h. **(E)** WT, DKO, TNFR1 TKO, ZBP1 TKO, and QKO B16 cells were pulsed with OVA peptide followed by co-culture with OT-I T cells at 1:4 E/T ratio. Target cell death was analyzed by IncuCyte. Values are triplicate mean ± SD. ****P<0.0001 by unpaired t test comparing last data point.

### TBK1 and IKKε restrains IFNγ-induced inflammatory apoptosis

To determine if other signaling pathways are affected by the loss of TBK1 and IKKε, we conducted global transcriptomics analysis which revealed that several pathways are induced by IFNγ to a greater degree in DKO cells compared to WT cells (Fig. 5A). One of the pathways enriched in IFNγ-stimulated DKO cells compared to WT cells comprise genes associated with the TNF-NFκB signaling pathway. Closer examination of the RNAseq data found enhanced expression of both canonical and non-canonical NFκB genes (e.g., *Rela*, *Relb*, *Nfkb1* and *Nfkb2*), as well as several NFκB-regulated inflammatory genes in IFNγ-stimulated DKO cells (Fig. 5B). We also conducted a global proteomics analysis which showed remarkable concordance with the transcriptomic analysis, including the enrichment of proteins associated with the TNF-NFκB signaling pathway in IFNγ-treated DKO cells (Fig. 5C). These omics analyses suggest that the NFκB pathway may be upregulated by IFNγ in the DKO cells, so we sought to study the functional relevance of these findings. To this end, nuclear lysate extracts were obtained from non-treated or IFNγ treated WT and DKO cells and examined by western blot. While little nuclear presence of NFκB proteins was detected in either control or IFNγ-stimulated WT samples, both canonical NFκB (RelA & p50) and non-canonical NFκB (RelB & p52) subunits were markedly elevated in the nuclear extracts from IFNγ treated DKO cells (Fig. 5D). Of note, complementation of DKO cells with TBK1 WT decreased nuclear translocation of NFκB subunits, while kinase-inactive TBK1 K38A cells did not (Extended Data Fig. 5A). These observations indicate that TBK1 prevents IFNγ from activating NFκB and reveal an unexpected regulation of the NFκB pathway by IFNγ. Since we showed earlier that apoptosis of the DKO cells induced by IFNγ was due to activation of TNFR1 and ZBP1, we asked if the two receptors also had a role in the hyperactivation of NFκB. Heightened expression of NFκB subunits in the DKO cells were not affected by the loss of ZBP1, whereas it was reduced by deletion of TNF or TNFR1 as seen in TNF TKO, TNFR1 TKO and QKO cells (Fig. 5D and Extended Data Fig. 5B). These results suggest that IFNγ stimulation of TBK1/IKKε-deficient cells leads to enhanced NFκB activation that is driven by TNFR1 signaling.

**Figure 5.**
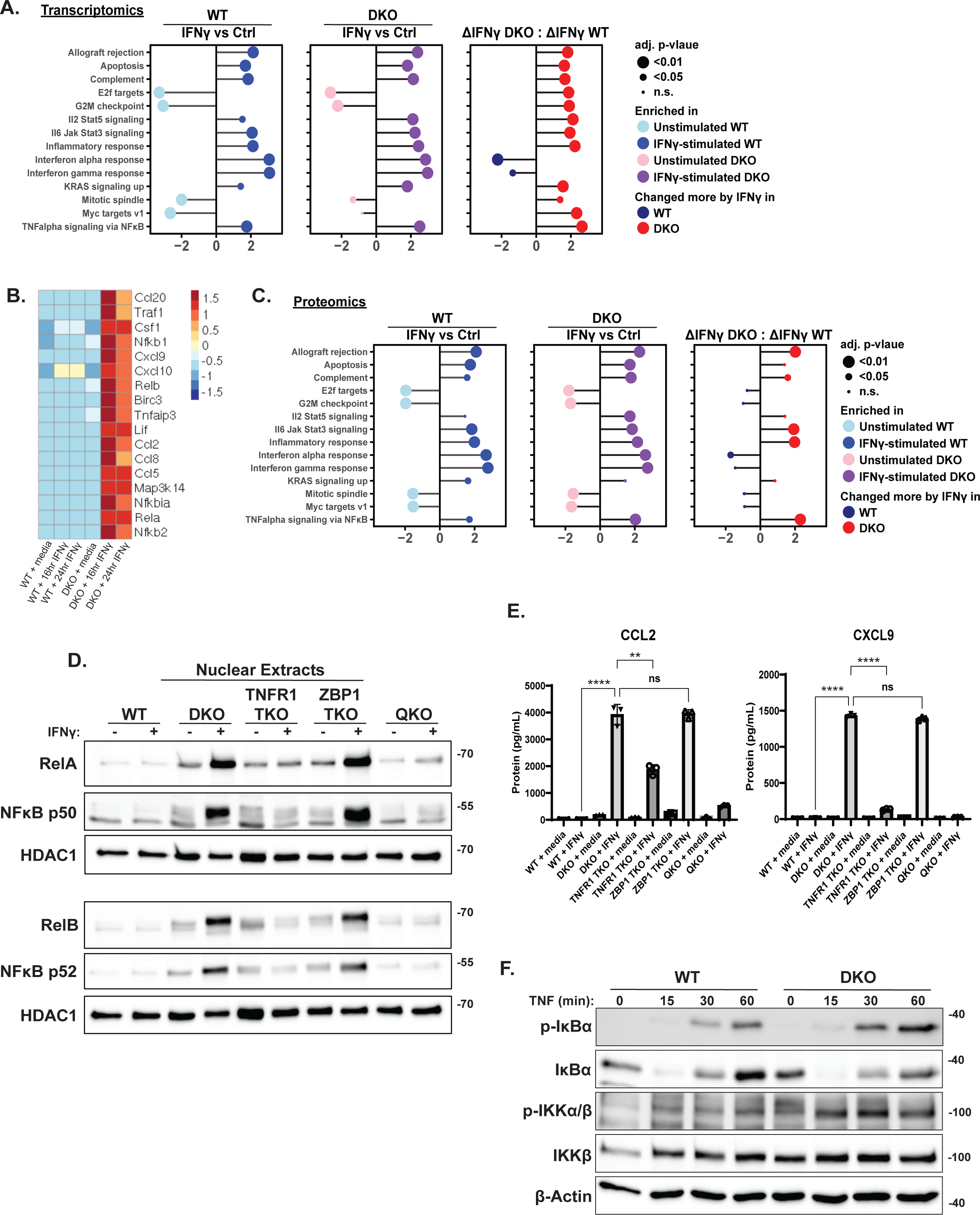
IFNγ induces inflammatory gene expression in TBK1/IKKε-deficient cells. **(A)** Global transcriptomics analysis of enriched gene pathways measured as log2 fold change between IFNγ**-** stimulated vs unstimulated control (Ctrl) WT cells (left), IFNγ**-**stimulated vs unstimulated control (Ctrl) DKO cells (middle), and a ratio of the IFNγ-stimulated change in DKO versus IFNγ-stimulated change in WT cells (right). **(B)** Heatmap expression profile of inflammatory genes obtained from RNAseq analysis of WT and DKO B16 cells stimulated with IFNγ (100 ng/mL) for 0, 16, 24 h. **(C)** Global proteomics analysis of enriched proteins in pathways measured as log2 fold change between IFNγ**-**stimulated vs unstimulated control (Ctrl) WT cells (left), IFNγ**-**stimulated vs unstimulated control (Ctrl) DKO cells (middle), and a ratio of the IFNγ-stimulated change in DKO versus IFNγ-stimulated change in WT cells (right). **(D)** Western blot analysis of the indicated proteins in nuclear extracts obtained from WT, DKO, TNFR1 TKO, ZBP1 TKO, and QKO B16 cells treated with IFNγ (100 ng/mL) for 0 or 24 h. **(E)** ELISA analysis of mouse CCL2 and CXCL9 in WT, DKO, TNFR1 TKO, ZBP1 TKO, and QKO B16 cells treated with IFNγ (100 ng/mL) for 0 or 24 h. ns = not significant, **P<0.01, ****P<0.0001, statistical analysis was performed using unpaired t test. **(F)** Western blot analysis of the indicated proteins in WT and DKO B16 cells treated with TNF (10 ng/mL) for 0, 15, 30, and 60 min.

To further examine the expression of inflammatory chemokines and cytokines observed in our transcriptomic and proteomic analyses, we performed a Luminex assay on culture supernatants harvested from non-treated or IFNγ treated WT and DKO cells at different timepoints. Consistent with the transcriptomic data, we detected significant increases in CXCL9, CCL2, LIF, and CSF1 only in supernatants obtained from stimulated DKO cells (Extended Data Fig. 5C). The upregulation of CXCL9 and CCL2 secretion in treated DKO cells was further validated by ELISA analysis (Fig. 5E). We also examined if CXCL9 and CCL2 upregulation is driven by TNFR1 and ZBP1. Similarly to the hyperactivation of NFκB, heightened chemokine expression was unaffected by the additional loss of ZBP1, whereas it was significantly reduced by the loss of TNFR1 as seen in TNFR1 TKO and QKO cells (Fig. 5E). These results strongly suggest that hyperactivation of NFκB and inflammatory gene expression in IFNγ stimulated TBK1/IKKε-deficient cells is driven by TNFR1 signaling.

Since IFNγ also induces death of TBK1/IKKε-deficient cells, we asked if the induction of the inflammatory gene program is a consequence of the RIPK1- and FADD-dependent death of these cells. Inhibiting death of the DKO cells with the RIPK1 kinase inhibitor Nec-1s, or by FADD deletion, did not affect the inflammatory gene program of NFκB and chemokine expression (Extended Data Fig. 5D-F). These results demonstrate that the induction of the inflammatory gene program in the DKO cells by IFNγ is independent of cell death and strongly suggests that TBK1/IKKε regulate another molecule other than RIPK1 to suppress NFκB. To identify this regulatory function of TBK1/IKKε, we first determined if the elevated inflammatory gene program was due to unrestrained NFκB. We tested this by transfecting an IκBα gene with S32A and S36A mutations, a non-degradable mutant commonly known as IκBα-super repressor (IκBSR), into the DKO cells (DKO-IκBSR) to inhibit NFκB nuclear translocation. The IκBSR inhibited the expression of CCL2 and CXCL9 (Extended Data Fig. 5F), consistent with the notion that NFκB is unrestrained in the DKO cells.

A previous study from Cohen and colleagues had suggested that TBK1/IKKε functions in a negative feedback manner by phosphorylating canonical IKKs (IKKα and IKKβ) to dampen their catalytic activity and NFκB signaling in innate signaling pathways^43^. We therefore postulated that TBK1/IKKε-deficient cells have a defect in this negative feedback control of IKKα/β, and the autocrine activation of TNFR1 after IFNγ stimulation would result in hyperactive NFκB. To test this possibility, we sought to see if there is elevated IKKα/β activity in the DKO cells. We initially attempted to analyze the phosphorylation of IκBα, the canonical IKK substrate, after IFNγ stimulation, but we were unable to obtain consistent results. This may be due to the slower kinetics of the response to IFNγ stimulation and the asynchronous nature of the autocrine activation of TNFR1. We reasoned that directly stimulating TNFR1 in a more synchronous manner with recombinant TNF would allow us to detect differences in IKK activity between the two cell lines. To this end, we treated WT and DKO cells with TNF and examined the phosphorylation of the IKKα/β kinase on Ser176/180, a target site for its auto-catalytic activity (Fig. 5F). There was more phospho-IKKα/β in the DKO cells compared to WT cells over 60 minutes, indicating enhanced IKKα/β activity. Note that the phospho-IKKα/β antibody is unable to distinguish between phosphorylated IKKα and IKKβ due to their homology. We also examined the phosphorylation of IκBα at Ser32/36, which was similarly elevated in the DKO cells. Once IκBα is phosphorylated, it undergoes degradation but is re-expressed in a feedback manner since it is a NFκB-inducible gene. This fluctuation in IκBα level is observed in the WT cells over the course of 60 minutes (Fig. 5F). However, in DKO cells, there is reduced IκBα protein at the later timepoints, indicative of continued phosphorylation and degradation of IκBα (Fig. 5F). These observations are consistent with a defect in negative feedback in the DKO cells, resulting in increased IKK and NFκB activity, providing a mechanistic explanation for the hyperactive expression of inflammatory genes. In sum, we showed that IFNγ can induce an inflammatory apoptotic death program when TBK1 and IKKε activity is compromised.

## Discussion

CTLs utilize perforin and members of the TNF superfamilies (TNFSF) to kill target cells, but it is unclear whether they can deploy other cytotoxic mechanisms. CTLs are also major producers of IFNγ and our study now describes how IFNγ can trigger apoptotic death of target cells, but this is inhibited by TBK1 and IKKε. Target cells that lack TBK1/IKKε are killed by IFNγ in an indirect manner via its upregulation of TNFR1 and ZBP1 to induce RIPK1-dependent apoptosis (Fig. 6). Under normal physiological conditions, TNFR1 ligation recruits RIPK1 to the intracellular domain of TNFR1, often referred to as complex I, where it is rapidly modified by cIAP1/2/TRAF2 and LUBAC E3 ligases that catalyze K63-linked and M1-linked ubiquitination, respectively^44^. M1-linked polyubiquitin chains act as a scaffold for the recruitment of NEMO and its associated kinases, IKKα and IKKβ. Additionally, NEMO engages with the adaptor protein TANK that brings both TBK1 and IKKε to complex I and collectively with IKKα/β phosphorylate RIPK1^12, 25, 45^. Others have shown these phosphorylation sites to be multiple residues, including Ser25, Thr189 and Ser321.^16, 25, 46^ These post-translational modifications of RIPK1 functions as an early cell death checkpoint to prevent RIPK1 from associating with the death-signaling complex often referred to as complex II^47^. Therefore, without TBK1/IKKε phosphorylating RIPK1, cells succumb to TNFR1-induced death. Our transcriptomic analysis indicated that TNF is induced by IFNγ in TBK1/IKKε-deficient cells and deletion of TNF diminished IFNγ-induced death to a similar extent as that of TNFR1 deletion. Interestingly, we did not detect significant levels of soluble TNF in the culture supernatants from IFNγ-stimulated DKO cells, suggesting that TNF may have remained membrane-bound in these cells. Furthermore, while soluble blocking antibody against TNF was able to reduce IFNγ-induced death of the DKO cells, it was much less effective compared to the genetic knockout of TNF. The differential effect of blocking anti-TNF versus genetic deletion of TNF suggests that cell-autonomous ligand-receptor interaction is activating downstream signaling. Cell-autonomous activation of TNFR1 has been previously reported in myeloid cells^33, 34^.

**Figure 6.**
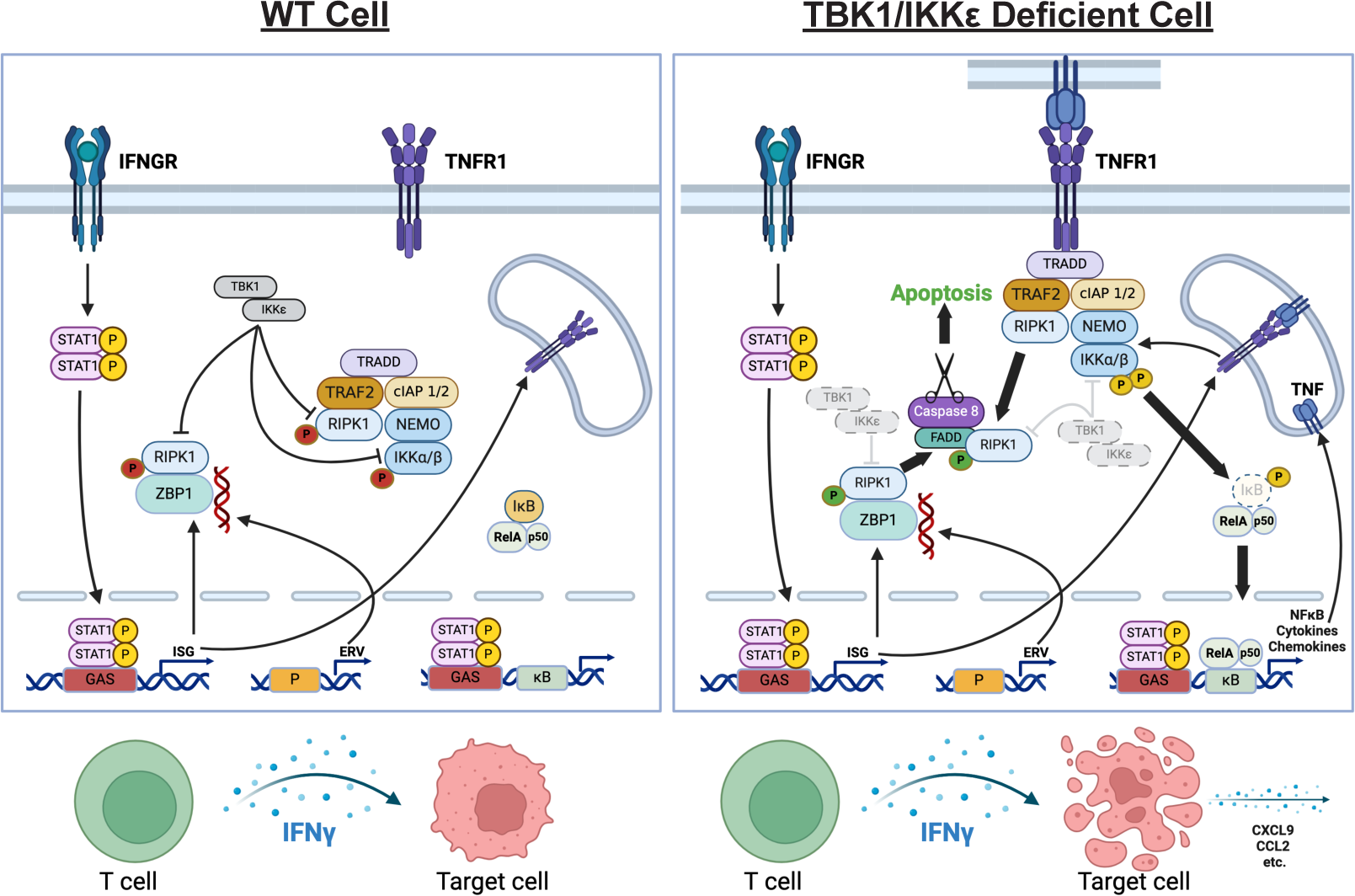
Model for the regulation of IFNγ-mediated death and inflammation by TBK1 and IKKε. Schematic of how multiple responses downstream of IFNγ stimulation are regulated by TBK1 and IKKε. (Left) In addition to their function in inducing type I IFN expression, TBK1 and IKKe also suppress RIPK1-dependent death and NFκB-driven inflammation. (Right) In their absence, IFNγ induces RIPK1-dependent apoptosis and NFκB-dependent inflammatory gene expression driven by autocrine activation of TNFR1. ZBP1 also plays a secondary role in these processes that is redundant to TNFR1. The figure was created using BioRender.

As TNF or TNFR1 deletion was unable to fully protect cells lacking TBK1/IKKε from IFNγ-induced cell death, we found that ZBP1 was the other signaling receptor contributing to the apoptotic death. ZBP1’s role in cell death has primarily been described as an activator of RIPK3-dependent necroptosis^40, 48^. In that response, RIPK1 inhibits the ability of ZBP1 to associate with RIPK3 to trigger necroptosis^49, 50^. The perinatal lethality of *Ripk1^-/-^* mice was shown to be due to excessive ZBP1-driven, RIPK3-dependent necroptosis^49, 50^. However, ZBP1-driven apoptosis has also been reported to occur in virally infected cells^51^. Since our B16 cells do not express RIPK3 nor do they appear to undergo necroptosis upon IFNγ stimulation, our data demonstrate that ZBP1 can also signal through RIPK1 to drive apoptosis in TBK1/IKKε-deficient cells. Nonetheless, TNFR1-driven apoptosis appears to be dominant as TNFR1 deletion has a more profound inhibitory effect on caspase activation in the DKO cells and was more protective against T cell killing, whereas ZBP1 deletion had minimal effect on caspase activation and was not protective against T cell killing.

The discovery that canonical and non-canonical NFκB and subsequent inflammatory chemokine expression was highly upregulated in only IFNγ stimulated DKO cells was unexpected for two reasons. One, it is widely accepted that NFκB has a pro-survival role. It was the first mechanism described to protect cells against death in the TNFR1 pathway via NFκB-dependent transcription of pro-survival genes such as cIAP1/2, TRAF2, c-FLIP and BCL2^11, 52, 53^. Thus, we did not expect NFκB activity to be present in cells that were also undergoing cell death, and our data suggest that the elevated NFκB activity in the DKO cells was insufficient to block RIPK1-dependent cell death. Two, the IFNγ receptor utilizes the JAK-STAT signaling mechanism and is not known to regulate NFκB. Our results now demonstrate that IFNγ stimulation can lead to NFκB-mediated gene transcription when TBK1/IKKε is absent, and this is manifested through the indirect activation of the TNFR1 pathway. Our analysis also indicates that the hyperactivation of NFκB occurs because TBK1/IKKε normally suppress the activity of IKKα/β. The juxtaposition of a NFκB inflammatory gene program with apoptosis in the absence of TBK1/IKKε reflects the dual function of the kinases in suppressing RIPK1-dependent death and IKK activity. This juxtaposition also argues against the notion that apoptotic death is non-inflammatory. Our observations suggest that under the appropriate context, apoptosis can be accompanied by inflammation. In this regard, a previous study using chimeric FKBP-RIPK1 reported that RIPK1-mediated death can be inflammatory^54^. Our study now demonstrates that IFNγ is a physiological signal that can induce inflammatory RIPK1-dependent apoptosis, but this is normally suppressed by TBK1/IKKε. It also suggests that the auto-inflammation observed in TBK1-deficient patients^17^ may be due to both inappropriate RIPK1-dependent death and an excessive NFκB-mediated inflammatory gene program.

Recently, Jenkins and colleagues reported that the deletion or inhibition of TBK1 in B16 melanoma tumors enhanced the efficiency of PD-1 blockade therapy^18^. In an in vivo CRISPR screen, they described *Tbk1* to be an immune-evasion gene. They reported that their TBK1-null B16 tumors were not sensitive to either TNF or IFNγ alone but were sensitive to a combination of the two cytokines. Our data provide evidence that either cytokine alone is sufficient to mediate cell death in tumor cells lacking TBK1/IKKε and furthermore, IFNγ produced by effector CD8 T cells is primarily responsible for killing TBK1/IKKε-deficient targets. We also uncovered the mechanistic basis for the sensitivity of TBK1/IKKε-deficient cells to IFNγ-induced RIPK1-dependent apoptosis and NFκB-dependent inflammatory response. While pharmacological inhibition of TBK1/IKKε in tumor cells has the potential to be useful in immunotherapy, the existence of the signaling circuitry we uncovered suggest an interesting evolutionary possibility. Amongst the molecules in the TNFR1 early cell death checkpoint that we tested, including NEMO, SHARPIN, TRAF2 and TBK1/IKKε, their deletion all confer sensitivity to TNF-induced apoptosis. However, only TBK1/IKKε deletion confers sensitivity to IFNγ-induced apoptosis, pointing towards a unique role for the two kinases. The results from this study place TBK1/IKKε at the center of three critical immune responses. The two kinases (1) induce type I IFN expression, (2) inhibit RIPK1-mediated cell death, and (3) inhibit NFκB-dependent inflammatory gene expression. We speculate that this functional wiring of TBK1 and IKKε is an evolutionary adaptation to pathogens that encode TBK1/IKKε inhibitors to block type I IFN expression as this would cause IFNγ to trigger infected cell demise, accompanied by an alternative inflammatory response driven by NFκB. This suggests that a disruption in an innate response may be compensated by an enhanced response to adaptive immunity. We further speculate that such pathogens would also need to encode IFNγ signaling antagonists to continue to exist.

## Materials & Methods

### Transduction of B16 F1 cell lines by lentivirus and retrovirus

For lentiviral VSVG pseudotyping, an 80% confluent 10 cm plate of HEK293 EBNA cells was transfected with 2.5 μg Peak8-VSVG, 7.5 μg psPAX2 encoding gag-pol (Addgene #12260), and 10 μg of lentiviral plasmid DNA packaged with 60 μL of Lipofectamine2000 (Thermo Fisher) in serum-free DMEM (Corning). The following day, the media was aspirated and replaced with fresh complete DMEM with 10% FBS, 100 IU/ml penicillin and 100 µg/mL streptomycin. After two days, the viral supernatants were harvested and concentrated in a Beckman Coulter Ultracentrifuge at 49,600 x g for 90 minutes at 4°C down to 1 mL. The viral supernatant was used to resuspend 1 x 10^6^ B16 F1 cells (provided by Miriam Merad, Icahn School of Medicine at Mount Sinai, New York, USA) with 4 μg/mL polybrene. The resuspended B16 cells were then plated in a six well plate, wrapped in saran wrap, and centrifuged at 859 x g for 90 minutes. Infected cells were cultured for three days after which the viral media was removed and replaced with fresh complete media with antibiotic drug for selection. Retroviral transduction was carried in a similar manner with the exception that pMD.OGP^55^ was used to express the retroviral gag-pol.

### Transduction of splenic T cells by retrovirus encoding OT-I TCR

For ecotropic retrovirus packaging, HEK293 EBNA cells was transfected with 5 μg pCL-Eco (provided by Dr. Larry Pease, Mayo Clinic) and 10 μg of TCR-2A-OTI-pMIG-II DNA (gift from Dr. Dario Vignali, Addgene #52111) as described above. The following day, the culture media was replaced with 10 mL of complete T cell media (RPMI 1640, 10% FBS, 100 IU/ml penicillin, 100 µg/mL streptomycin, 100 µM nonessential amino acids, 2 mM L-glutamine, 1 mM sodium pyruvate, and 50 µM of 2-mercaptoethanol) and cultured for another 24 h. The viral supernatant was harvested, centrifuged at 483 x g for 5 minutes to pellet cellular debris. The top 8 mL of viral supernatant was collected and used to resuspend 10^6^ *Prf1^+/+^* or *Prf1^-/-^* splenic cells with 4 μg/mL polybrene and 50 U/mL of IL-2. The resuspended T cells were plated in a 24-well plate pre-treated with RetroNectin and centrifuged for 90 min at 800 x g at 32°C. The cells were then given 1 mL of additional T cell media with Con A (2.5 ug/mL) and IL-2 (50 U/mL) and cultured in a 37°C incubator for 1 day. T cells were then split into a new 24-well plate and incubated for an additional 2 days prior to use.

### Generation of B16 F1 knockout cell lines

Compound knockouts of B16 cells were generated by sequential lentivirus transduction using pLenticrispr v2 (gift from Dr. Feng Zhang, Addgene #52961, puromycin selection), lenti-sgRNA blast (gift from Dr. Brett Springer, Addgene #104993, blasticidin selection), lenti-sgRNA neo (gift from Dr. Brett Springer, Addgene #104992, G418 selection) and lenti-sgRNA hygro (gift from Dr. Brett Springer, Addgene #104991, hygromycin selection). At each knockout and antibiotic-selection stage, a corresponding non-targeting control guide was introduced using the same vector. Lentiviral plasmids expressing sgRNA were either purchased from Genscript or generated in-house using oligonucleotides synthesized by Integrated DNA Technologies (IDT).

### Lentiviral guide RNA target sequences for generating knockouts

NT (non-targeting): GCGAGGTATTCGGCTCCGCG

*Tbk1:* CAACATCATGCGCGTCATAG

*Ikbke:* CATCGTGAAGCTATTCGCAG

*Tnfrs1a:* GTGTCTCACTCAGGTAGCGT

*Tnf:* GTAGACAAGGTACAACCCAT

*Zbp1:* AGTCCTTTACCGCCTGAAGA

*Fadd:* CCGCAGCGCCTTAACCAGTC

*Casp8:* TGAGATCCCCAAATGTAAGC

*Stat1:* GGTCGCAAACGAGACATCAT

### Retroviral/lentiviral expression constructs

The RetroHygro vector used for stable expression was generated in-house from the Moloney murine leukemia virus (MMLV) vector pMMP412^56^ into which an internal ribosome entry site (IRES)-hygromycin resistance cassette was inserted downstream of the ORF of interest. ORFs encoding FLAG-tagged human *TBK1*, *TBK1-K38A*, and *IKBKE* were amplified from existing plasmids^17^ and cloned into RetroHygro by conventional or In-Fusion technique. The ORF encoding Strep II-tagged p150 isoform of murine ADAR1 was synthesized by Twistbio and cloned into RetroHygro. The lentiviral plasmid for expressing cytoplasmic ovalbumin (cOVA) expression was generated by cloning gene fragments synthesized by Twistbio into the pLVX-EF1-alpha vector. The ORF encodes a cOVA-T2A-Hygro-P2A-GFP-NLS polypeptide.

### Generation of knockout SVEC4-10 cell line by ribonucleoprotein (RNP) transfection

For each gene, two different target crRNAs (100 μM) were individually combined with tracrRNA (100 μM) at a 1:1 ratio and incubated at 95°C for 5 minutes to form duplexes. crRNAs and tracrRNA were synthesized by IDT. A mixture of 4.5 μL of each duplex and 4.5 μL of Cas9 enzyme (MacroLab Facility, UC Berkeley) was incubated at room temperature for 10 minutes to form RNP complex. 1 million SVEC4-10^57^ cells (provided by Dr. Douglas Green, St Jude’s Children’s Research Hospital, Tennessee, USA) were resuspended in Lonza SF nucleofection buffer and combined with the RNP complex. Cells were electroporated using Lonza 4-D nucleofector pulse code DJ-110. Knockouts were validated 48 hours post electroporation by western blot. The two guide target sequences for *Tbk1* are ACGGGGCTACCGTTGATCTG and TTTGAACATCCACTGGGCGA.

### Isolation of OT-I T cells

6-well tissue culture plates were coated overnight at 4°C with an anti-CD3 antibody (Bio X Cell) in PBS. The following day, spleens from 8-12 weeks old OT-I mice (Jax #3831) were mashed and filtered through a 40 μm filter. The filtrate is centrifuged and red blood cells lysed with 1 mL ACK lysis buffer for 5 minutes. PBS was then added to dilute the lysis buffer followed by centrifugation. After removing the supernatant, T cells were purified using the EasySep Mouse Pan-Naïve T Cell Isolation Kit (STEMCELL). T cells were then cultured for 24 hours in coated anti-CD3 plate with the addition of anti-CD28 and IL-2 to activate T cells.

### Animals and tumor challenges

All animal studies were carried out in accordance with approved IACUC protocol at the Mayo Clinic in Rochester, MN. C57BL/6 (Jax #664) and C57BL/6-Tg (TcraTcrb) 1100Mjb/J (OT-I) (Jax #3831) were obtained from Jackson Laboratory and housed in standard mice rooms. NOD.Cg-Prkdc^scid^il2rg^tm1Wjl^/SzJ (NSG) (Jax #5557) were obtained from Jackson Laboratory and housed in barrier mice rooms. Tumor challenge experiments were performed with mice 8 weeks or older. Mice were shaved at the inoculation site a day before tumor implantation. 0.1 x 10^6^/100 μL B16 F1-cOVA tumor cells were resuspended in PBS (Corning) and subcutaneously injected into the right flank on day 0. When all tumors became palpable (day 12), tumor volume was measured with a digital caliper and randomized for the single treatment of either PBS (100 μL) or isolated OT-I T cells (10 x 10^6^/100 μL) through intravenous injection via tail vein. Tumor volume was measured every 2-3 days until either survival end point was reached or till the end of study on day 42. Tumor end points were adhered to as defined by the IACUC protocol. Mice were euthanized by AVMA-approved CO_2_ asphyxiation. At least five mice were used in each group for all experiments.

### Western blotting

Whole-cell lysates were obtained using triton lysis buffer (20 mM Tris-HCl pH 7.4, 40 mM NaCl, 5 mM EDTA, 50 mM NaF, 30 mM Na Pyrophosphate, 1% Triton X-100) that contained 1X protease inhibitor (Millipore Sigma, 539137) and 1X phosphatase inhibitor (Thermo Scientific, 78426). Protein concentration was measured using Pierce BCA (Thermo Scientific, 23227). 50 μg of protein samples were boiled at 95°C in 1X SDS sample buffer and resolved by reducing SDS-PAGE. Resolved proteins were transferred to nitrocellulose membranes (Amersham, 10600003), blocked with 5% milk in 1X TBST solution for 1 h at room temperature, followed by overnight incubation with primary antibodies at 4°C. After a series of washes with 1X TBST, membranes were incubated with secondary HRP antibodies in 2.5% milk in 1X TBST solution. After multiples washes, membranes were incubated in chemiluminescent substrate solution (Thermo Scientific, 34076) for 2 minutes and imaged with the BIO-RAD ChemiDoc MP instrument. For reblotting, membranes were stripped with guanidine HCl prior, blocked with milk and re-probed with subsequent antibody.

### Co-immunoprecipitation

For each condition, two confluent 10 cm plates of tumor cells were stimulated and then lysed in buffer containing 30 mM Tris-HCl (pH7.4), 150 mM NaCl, 10% glycerol, 1% Triton X-100, 30 mM Na Pyrophosphate, 50 mM NaF, protease inhibitor, and phosphatase inhibitor. Lysates were cleared by centrifugation at 18,407 x g at 4°C, and protein concentration was measured using Pierce BCA (Thermo Scientific). A fraction of the lysates was set aside for analyzing total protein analysis. For ZBP1 immunoprecipitation, an equivalent amount of protein in each sample was immunoprecipitated by rotating with 5 ng ZBP1 mAb (Adipogen, AG-20B-0010-C100) overnight at 4°C. Immune complexes were affinity purified with Protein A/G beads. After extensive washes, the beads were eluted with SDS-sample buffer at 70°C for 20 minutes. Immunoprecipitated protein complexes and 50 μg of total lysates were resolved by SDS-PAGE and sequentially blotted with primary antibodies.

### Nuclear lysate extractions

After stimulation, cell cultures were counted and an equivalent number of cells from each sample were resuspended in 400 µL of Buffer A containing 10 mM HEPES (pH 7.9), 10 mM potassium chloride, 0.1 mM EDTA, 0.1 mM EGTA, protease inhibitor, and phosphatase inhibitor. After incubation on ice for 15 minutes, 25 μL of 10% NP-40 was added to samples and vigorously vortexed. Samples were then centrifuged at 18,407 x g for 1 minute and supernatants were collected as cellular extract. The pellet was washed once with Buffer A and resuspended in a buffer containing 20 mM HEPES (pH 7.9), 0.4 M sodium chloride, 1 mM EDTA, 1mM EGTA, protease inhibitor, and phosphatase inhibitor. Samples were then shaken for 30 minutes at 4°C, centrifuged at 18,407 x g for 1 minute, and the supernatants were collected as nuclear lysates. Lysates were then resolved by SDS-PAGE for western blotting.

### Cell death quantification

Target tumor cells were seeded at 4000 cells/well in a 96-well tissue culture plate. After overnight culture, the old media was replaced with fresh complete media containing recombinant cytokines or purified OT-I T cells together with 0.5 μM of the cell-impermeable viability dye YOYO-3 (Thermo Scientific, Y3606). The cultures were analyzed using a Sartorius IncuCyte S3 live-cell imaging system and four images of each well were taken every 4 hours. Cell death events were quantified as a measure of YOYO-3 fluorescence counts normalized to the confluency at each time point. Data shown are the mean of triplicate samples ± SD and are representative of at least 3 replicated experiments.

### ELISA

Cells were seeded at 0.125 x 10^6^ cells/well in a 6-well tissue culture plate. After overnight culture, old media was replaced with fresh complete culture media containing either recombinant cytokines or purified OT-I T cells. After treatment for the indicated times, supernatants were collected and centrifuge to remove cellular debris. Supernatants were then tested for cytokine or chemokine detection by ELISA. The ELISA kits were obtained from the following sources: mouse TNFα (BioLegend, 430904), mouse IFNγ (BioLegend, 430801), mouse MCP-1 (CCL2) (BioLegend, 432704), and mouse MIG (CXCL9) (Bio-Techne, DY492-05).

### RNA isolation and RT-qPCR

Cells were seeded in 6-well plates and cultured overnight at 37°C. They were then treated with either media or 100 ng/mL IFNγ for 24 hr. Total RNA was extracted using the RNeasy Mini Kit (Qiagen, 74104) according to manufacturer’s protocol and concentration measured using a nanodrop (Thermo Fisher NanoDrop One). Equal amounts of RNA were used for reverse transcription to generate cDNA. cDNA, SYBR Green SuperMix (Quantabio, 95056-500) and primers were mixed and subjected to qPCR using the BIO-RAD CFX Connect Real-Time System. The following primers were used: *Tnf*, forward 5’-TATGGCTCAGGGTCCAACTC-3’ and reverse 5’-CTCCCTTTGCAGAACTCAGG-3’; *Gapdh,* forward 5’-AACGACCCCTTCATTGAC-3’ and reverse 5’-TCCACGACATACTCAGCAC-3’. *Tnf* mRNA was normalized to *Gapdh*.

### Bulk RNA-seq and analysis

B16 tumor cells were seeded in 6-well plates and cultured overnight at 37°C. The cells were then treated with either media or 100 ng/mL IFNγ for 16 and 24 hr. Total RNA was extracted from tumor cells using the RNeasy Mini Kit (Qiagen, 74104) according to manufacturer’s protocol. RNA samples were shipped to BGI Genomics for bulk RNA-seq on the DNBseq platform, yielding 20 million high quality paired-end 100 bp reads with ≥80% bases with Q30 score. High quality reads were aligned to the mouse reference genome build GRCm39 and a transcript expression count matrix was generated using Rsubread^58^. Read counts were normalized for library size using Limma-voom and analyzed for differential expression using the “limma” R package^59^.

### Sample preparation for mass spectrometry

B16 F1 cells were treated with media or IFNγ for 24 h in biological triplicates. After stimulation, cells were lysed in lysis buffer (9 M urea, 50 mM triethylammonium bicarbonate (TEABC), 1 mM sodium orthovanadate, 1 mM β-glycerophosphate, 2.5 mM sodium pyrophosphate) by sonication. After centrifugation, the protein extracts from each sample were reduced, alkylated, and subjected to in-solution trypsin digestion. After desalting and lyophilization, the resulting peptides from each sample were labeled with a TMTpro reagent, separately. The TMT-labeled peptides were mixed and fractionated by basic pH reversed-phase chromatography (bRPLC) to 96 fractions, which then were concatenated to 12 fractions. An aliquot (5%) of each fraction of peptides were subjected to LC-MS/MS analysis for global proteomic analysis.

### LC-MS/MS analysis

Peptides from each fraction were analyzed by LC-MS/MS on an Orbitrap Exploris 480 mass spectrometer (ThermoFisher, Bremen, Germany) using a gradient 150-minute LC method. Data-dependent acquisition (DDA) was set as: MS1 survey scan data from m/z 340-1800 at 120,000 resolution (at m/z 120), 300% AGC, max fill time of 100 ms; MS/MS scan from m/z 110 at 45,000 resolution, a minimum precursor intensity of 70,000, quadrupole isolation width of 0.7 Thompson, 100% AGC target, max fill time of 120 ms, NCE=33, for precursor charge states of 2-4.

### MS data analysis

The mass spectra were searched against a UniProt mouse protein database (2024_02 version) by Andromeda algorithm on the MaxQuant (ver. 2.2.0.0) proteomics analysis platform. The search parameters were set: carbamidomethylation on cysteine residues, TMTpro-16plex modification on N-terminal and lysine residues as fixed modifications; protein N-terminal acetylation, oxidation on methionine residues; a maximum of two missed cleavages. The data were searched against target decoy database and the false discovery rate was set to 1% at the peptide level. The quantile-normalized and log-transformed reported ion intensities were used to quantitate the changes of proteins between different conditions^59^.

### Pathway enrichment analysis

We performed Gene Set Enrichment Analysis (GSEA) using the “fgsea” R package^60^. Briefly, the genes were firstly ranked by log-transformed and signed *P*-values obtained from transcriptomic or proteomic comparisons between unstimulated control (Ctrl) vs IFNγ-stimulated WT cells, unstimulated control (Ctrl) vs IFNγ-stimulated DKO cells, and the IFNγ-stimulated change in DKO versus IFNγ-stimulated change in WT cells, and then analyzed against the mouse hallmark gene set collection from the MSigDB database. The GSEA results were visualized in R using ggplot2 package^61^.

### Luminex Assay

Cells were seeded at 2 x 10^6^ cells in a 10 cm tissue culture plate. The next day, the media was replaced with fresh complete culture media containing recombinant cytokines. Supernatants were harvested at indicated times and spun down to remove any cellular debris. Supernatants were then tested for cytokine or chemokine detection with a custom designed ProcartaPlex Luminex panel from ThermoFisher that includes mouse MCP-1/CCL2 (EPX01A-26005-901), MIG/CXCL9 (EPX010-26061-901), LIF (EPX01A-26040-901), and CSF1 (EPX01A-26039-901).

### Reagents and Antibodies

Cytokines and reagents used were from the following sources: TNF (PeproTech, 315-01A), IFNγ (PeproTech, 315-05), IFNβ (Bio-Techne, 8234-MB-010), Necrostatin-1s (MedChemExpress, HY-14622A), Ruxolitinib (MedChemExpress, 50-202-9341) LCMV GP33 (GenScript, RP20257), and OVA peptide (257-264) (GenScript, RP10611). Antagonist antibodies used were from following sources: TNF (clone XT3.11, Bio X Cell BE0058) and IFNγ (clone XMG1.2, Bio X Cell BE0055). For western blotting, primary antibodies used were TBK1 (Cell Signaling, 3013S), IKKε (Cell Signaling, 2690S), phospho-IRF-3 (Ser396) (Cell Signaling, 4947S), IRF-3 (Cell Signaling, 4302S), phospho-STAT1 (Tyr701) (Cell Signaling, 9167S), STAT1 (Cell Signaling, 9172S), phospho-RIPK1 (Ser166) (Cell Signaling, 31122S), RIPK1 (Cell Signaling, 3493S), cleaved caspase 8 (Cell Signaling, 8592S), cleaved caspase 3 (Cell Signaling, 9661S), cleaved PARP (Cell Signaling, 9541S), FADD (Abcam, ab124812), RIPK3 (ProSci, 2283), phospho-MLKL (Ser345) (Cell Signaling, 62233S), MLKL (Millipore Sigma, MABC604), TNFR1 (Cell Signaling, 13377S), ZBP1 (Adipogen, AG-20B-0010-C100), Strep II tag (GenScript, A01732), NFκB p65/RelA (Cell Signaling, 8242S), NFκB p105/p50 (Cell Signaling, 12540S), NFκB p100/p52 (Cell Signaling, 52583S), RelB (Cell Signaling, 4922S), phospho-IκBα (Ser32/36) (Cell Signaling, 9246S), IκBα (Cell Signaling, 4812S), phospho-IKKα/β (Ser176/180) (Cell Signaling, 2694S), and IKKβ (Cell Signaling, 8943S). β-Actin (Cell Signaling, 3700S) was used as a loading control for whole cell lysates, while HDAC1 (Cell Signaling, 5356T) was used as a loading control for nuclear lysates. Primary antibodies were used at a 1:1000 dilution in antibody buffer containing 2.5% BSA, 0.05% sodium azide in 1X TBST. Secondary antibodies against rabbit IgG (Jackson ImmunoResearch, 111-035-144), mouse IgG (Jackson ImmunoResearch, 115-035-146), and rat IgG (Jackson ImmunoResearch, 112-035-143) were used at 1:5000 in 2.5% milk in 1X TBST.

### Statistics

We performed statistical analysis using Prism GraphPad software version 10. For cell death data produced by the IncuCyte, we used 2-tailed student’s t-test with normal distribution comparing the last time point between the two groups. 2-tailed student’s t-test was implemented comparing two groups to each other in ELISA and Luminex data. Ordinary 1-way ANOVA test was used when comparing multiple experimental groups. For mice tumor volume data, we implemented 2-tailed paired t-test to compare the two groups. P < 0.05 was considered significant.

## Author contributions

N.D.S., Y.C., and E.N.K. designed and performed the experiments. A.R.C., J.Z., M.C.G., M.C., and M.A.H. developed reagents and conducted the experiments. H.D., A.P., and L.M.R. provided technical guidance, analyzed the data, and edited the manuscript. D.B. provided reagents and edited the manuscript. N.D.S., A.R.C., J.Z., and A.T.T. wrote the manuscript. A.T.T. directed the studies.

## Acknowledgements

We thank members of the Ting lab for helpful discussions. We thank Dr. Aaron Johnson (Mayo Clinic) for providing the *Prf1^-/-^* mice. This work was supported by the Mayo Clinic and by National Institutes of Health (NIH) grants AI052417 (A.T.T), CA270380 (A.T.T & H.D.), AI148963 (A.T.T. & D.B.). N.D.S., M.C.G and M.A.H. were supported by a T32 Training Program in Immunology AI170478. We also thank the assistance of the Mayo Clinic Proteomics Core, which is a shared resource of the Mayo Clinic Comprehensive Cancer Center (NCI P30 CA15083).

