## Extended Data for "TBK1 and IKKε protect target cells from IFNγ-mediated T cell killing via an inflammatory apoptotic mechanism"

### Extended Data Figure 1

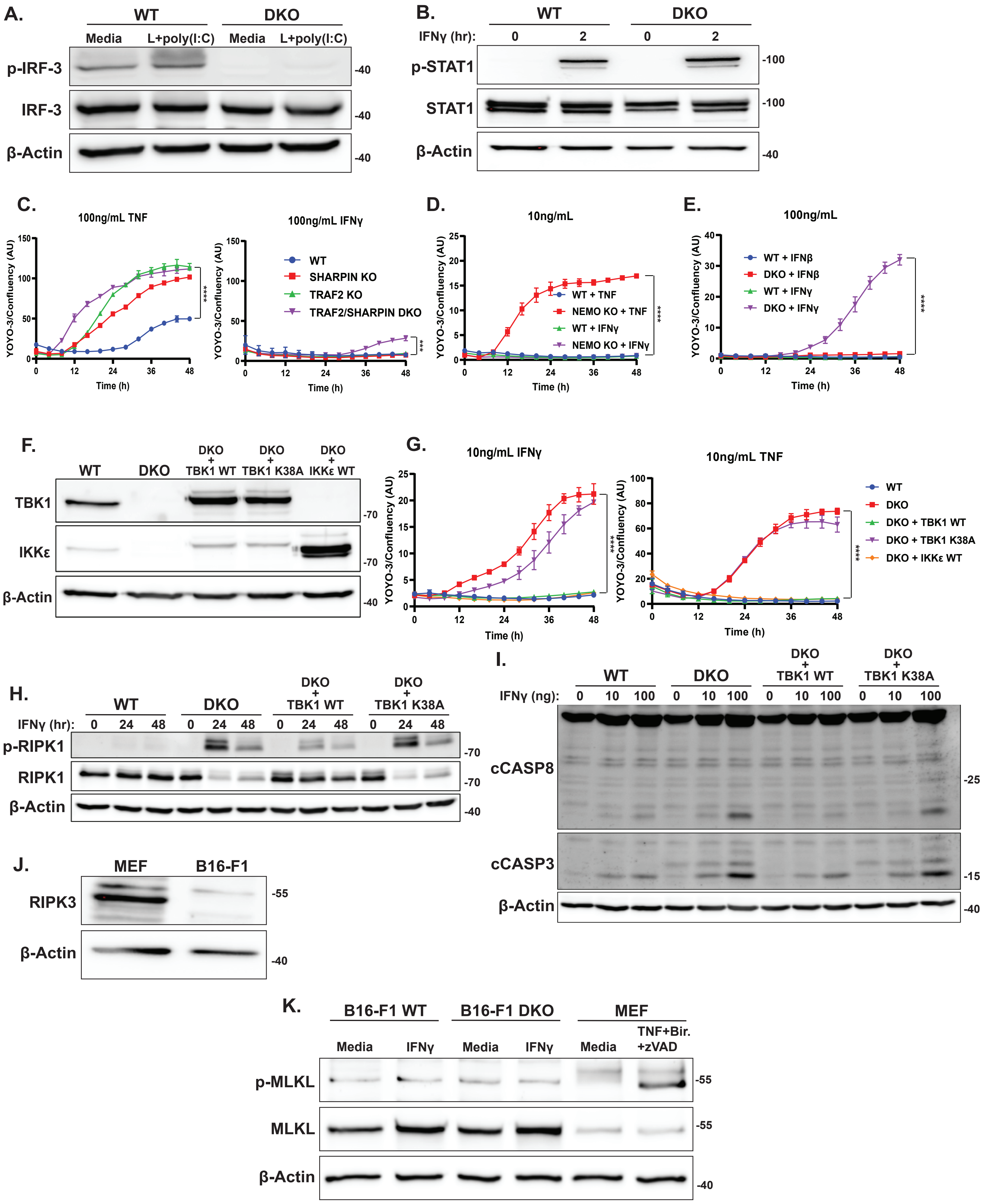

**Extended Data Figure 1. Supporting data for the kinase-dependent role of TBK1 in suppressing IFNγ-mediated apoptosis. (A)** Western blot analysis of the indicated proteins in WT and DKO B16 cells treated with media alone or lipofectamine 2000 (10 μL) + poly I:C (1 μg) for 8 h. **(B)** Western blot analysis of the indicated proteins in WT and DKO B16 cells treated with IFNγ (100 ng/mL) for 0 and 2 h. **(C)** WT, Sharpin KO, TRAF2 KO, and Sharpin/TRAF2 DKO B16 cells were treated with TNF (100 ng/mL; left) or IFNγ (100 ng/mL; right) and analyzed by IncuCyte. Values are triplicate mean ± SD. ***P<0.001, ****P<0.0001 by unpaired t test comparing last data point. **(D)** WT and NEMO KO B16 cells treated with TNF (10 ng/mL) or IFNγ (10 ng/mL) and analyzed by IncuCyte. Values are triplicate mean ± SD. ****P<0.0001 by unpaired t test comparing last data point. **(E)** WT and DKO B16 cells treated with IFNβ (100 ng/mL) or IFNγ (100 ng/mL) and analyzed by IncuCyte. Values are triplicate mean ± SD. ****P<0.0001 by unpaired t test comparing last data point. **(F)** Western blot analysis of the indicated proteins in WT, DKO, DKO + TBK1 WT, DKO + TBK1 K38A, DKO + IKKε WT B16 cells. **(G)** WT, DKO, DKO + TBK1 WT, DKO + TBK1 K38A, DKO + IKKε WT B16 cells were treated with IFNγ (10 ng/mL; left) or TNF (10 ng/mL; right) and analyzed by IncuCyte. Values are triplicate mean ± SD. ****P<0.0001 by unpaired t test comparing last data point. **(H)** Western blot analysis of the indicated proteins in WT, DKO, DKO + TBK1 WT, and DKO + TBK1 K38A B16 cells pretreated with zVAD-fmk (20 μM) for 30 min followed by IFNγ (100 ng/mL) for 0, 24, 48 h. **(I)** Western blot analysis of the indicated proteins in WT, DKO, DKO + TBK1 WT, and DKO + TBK1 K38A B16 cells treated with 0, 10, or 100 ng/mL IFNγ for 24 h. **(J)** Western blot analysis of the indicated proteins in mouse embryonic fibroblasts (MEF) and B16 cells. **(K)** Western blot analysis of the indicated proteins in WT and DKO B16 cells treated with media or IFNγ (100 ng/mL) for 24 h, and MEF cells treated with media or the combination of TNF (100 ng/mL) + Birinapant (10 μM) + zVAD-fmk (20 μM) for 2 h.

### Extended Data Figure 2

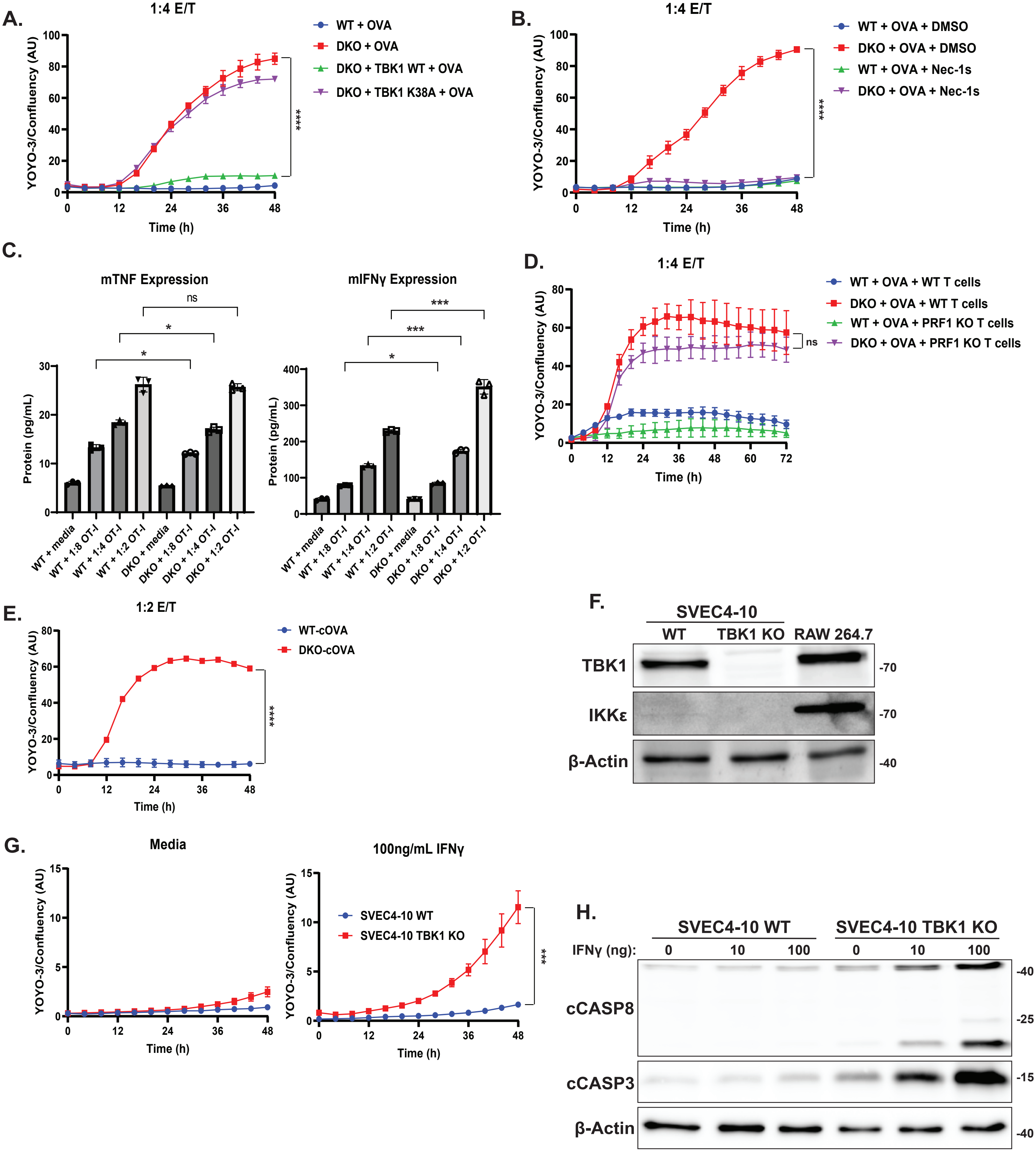

**Extended Data Figure 2. Supporting data for T cell killing of TBK1/IKKε-deficient target cells and characterization of IFNγ sensitivity in SVEC4-10 cells. (A)** WT, DKO, DKO + TBK1 WT, and DKO + TBK1 K38A B16 cells were pulsed with OVA peptide followed by co-culture with OT-I T cells at 1:4 E/T ratio. Target cell death was analyzed by IncuCyte. Values are triplicate mean ± SD. ****P<0.0001 by unpaired t test comparing last data point. **(B)** WT and DKO B16 cells were pulsed with OVA peptide followed by co-culture with OT-I T cells at 1:4 E/T ratio in the presence of DMSO or Nec-1s (10 μM). Target cell death was analyzed by IncuCyte. Values are triplicate mean ± SD. ****P<0.0001 by unpaired t test comparing last data point. **(C)** ELISA analysis of mouse TNF (left) or IFNγ (right) in WT and DKO cells co-cultured with none or OT-I T cells at 1:8, 1:4, 1:2 E/T ratio for 24 h. ns = not significant, *P<0.05, ***P<0.001, statistical analysis was performed using unpaired t test. **(D)** WT and DKO B16 cells were pulsed with OVA peptide followed by co-culture with WT or Perforin (Prf1) KO T cells expressing OT-I TCR at 1:4 E/T ratio. Target cell death was analyzed by IncuCyte. Values are triplicate mean ± SD. ns = not significant, by unpaired t test comparing last data point. **(E)** WT-cOVA and DKO-cOVA B16 cells were co-cultured with OT-I T cells at 1:2 E/T ratio. Target cell death was analyzed by IncuCyte. Values are triplicate mean ± SD. ****P<0.0001 by unpaired t test comparing last data point. **(F)** Western blot analysis of the indicated proteins in WT and TBK1 KO SVEC4-10 tumor cells and RAW264.7 macrophage cells as positive control. Knockout in SVEC4-10 cells were generated by nucleoporation of Cas9-sgRNA ribonucleoprotein (RNP) complexes. **(G)** WT and TBK1 KO SVEC4-10 cells were treated with media alone or IFNγ (100 ng/mL) and analyzed by IncuCyte. Values are triplicate mean ± SD. ***P<0.001 by unpaired t test comparing last data point. **(H)** Western blot analysis of the indicated proteins in WT and TBK1 KO SVEC4-10 cells treated with 0, 10, or 100 ng/mL IFNγ for 24 h.

### Extended Data Figure 3

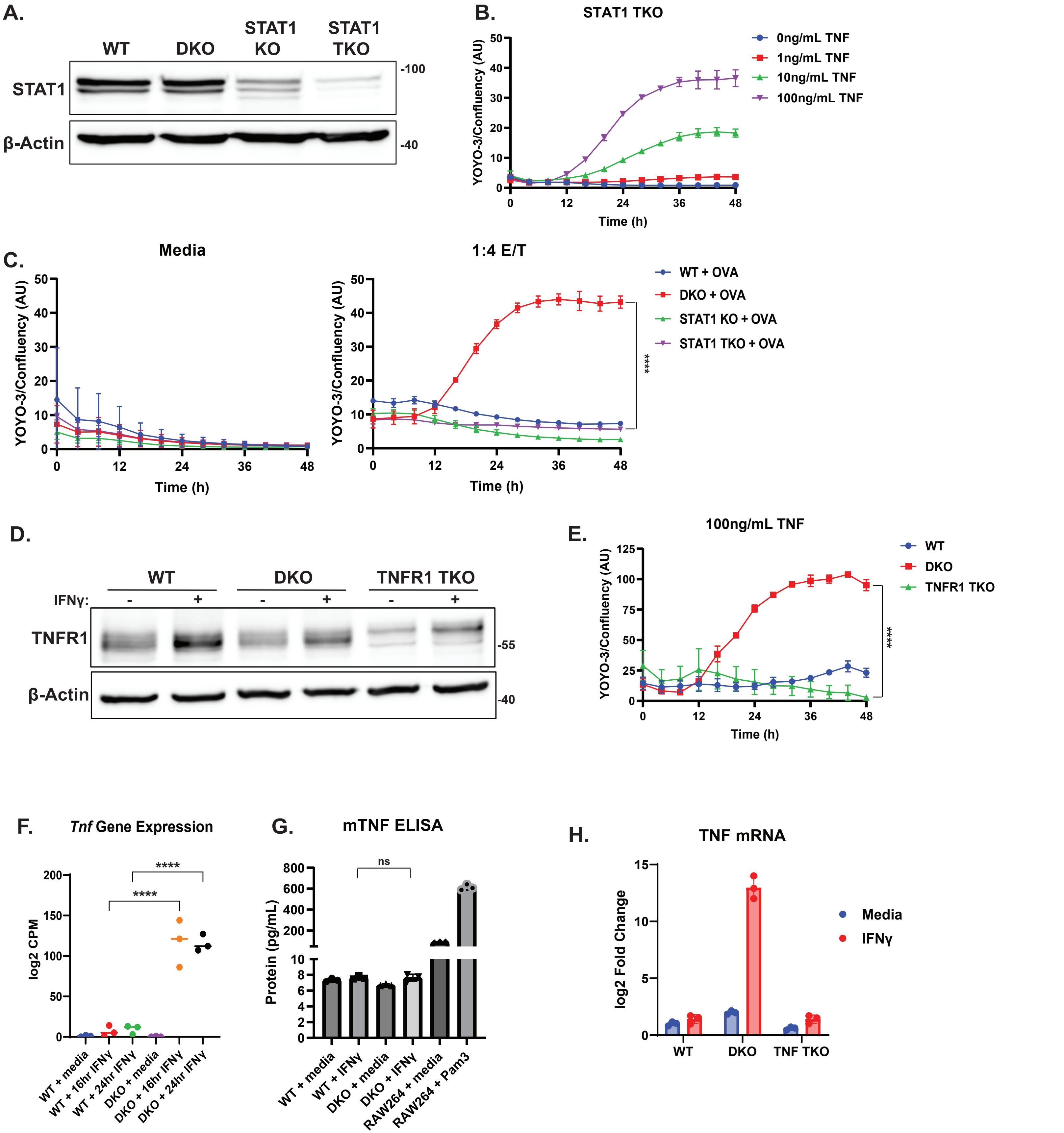

**Extended Data Figure 3. Supporting data for the role of TNF and TNFR1 in IFNγ-mediated killing of TBK1/IKKε-deficient cells. (A)** Western blot analysis of the indicated proteins in WT, DKO, STAT1 single KO, and TBK1/IKKε/STAT1 TKO B16 cells. **(B)** TBK1/IKKε/STAT1 TKO B16 cells treated with 0, 1, 10, 100 ng/mL TNF and analyzed by IncuCyte. Values are triplicate mean ± SD. **(C)** WT, DKO, STAT1 KO, and TBK1/IKKε/STAT1 TKO B16 cells were pulsed with OVA peptide followed by co-culture with none (left) or OT-I T cells at 1:4 E/T ratio (right). Target cell death was analyzed by IncuCyte. Values are triplicate mean ± SD. ****P<0.0001 by unpaired t test comparing last data point. **(D)** Western blot analysis of the indicated proteins in WT, DKO, and TBK1/IKKε/TNFR1 TKO B16 cells treated with media or IFNγ (100 ng/mL) for 24 h. **(E)** WT, DKO, TBK1/IKKε/TNFR1 TKO B16 cells treated with TNF (100 ng/mL) and analyzed by IncuCyte. Values are triplicate mean ± SD. ****P<0.0001 by unpaired t test comparing last data point. **(F)** WT and DKO B16 cells were stimulated with IFNγ (100 ng/mL) for 0, 16, 24 h and RNA isolated for sequencing. Values displayed as log2 CPM comparing *Tnf* from three independent experiments. Statistical analysis was performed using one-way ANOVA with Sidak's multiple-comparison test. ****P<0.0001. **(G)** ELISA analysis of mouse TNF in supernatants from WT and DKO B16 cells treated with media or IFNγ (100 ng/mL) for 24 h. RAW264 cells treated with media or Pam3CSK4 (1 ng/mL) was used as positive control. ns = not significant, statistical analysis was performed using unpaired t test. **(H)** RT-qPCR analysis of TNF mRNA from WT, DKO, and TBK1/IKKε/TNF TKO B16 cells treated with media or IFNγ (100 ng/mL) for 24 h.

### Extended Data Figure 4

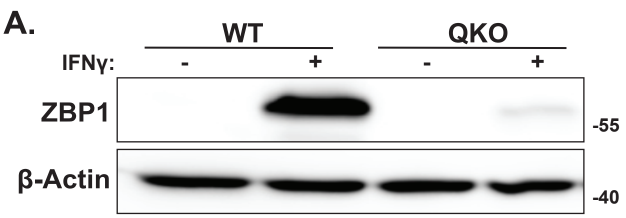


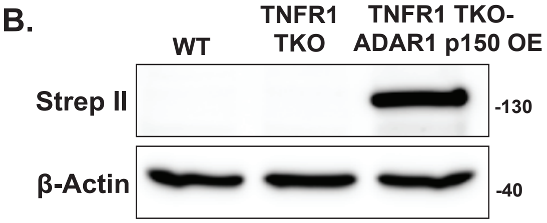


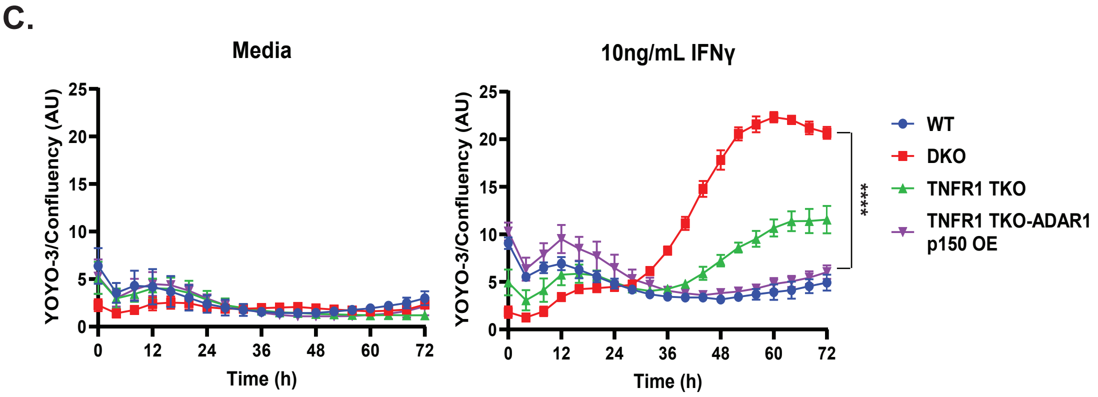

**Extended Data Figure 4. Supporting data for the co-involvement of ZBP1 and TNFR1 in mediating IFNγ killing of TBK1/IKKε-deficient cells. (A)** Western blot analysis of the indicated proteins in WT or TBK1/IKKε/TNFR1/ZBP1 QKO B16 cells treated with media or IFNγ (100 ng/mL) for 24 h. **(B)** Western blot analysis of the indicated proteins in WT, TNFR1 TKO, and TNFR1 TKO-ADAR1 p150 OE (Strep II tagged) B16 cells. **(C)** WT, DKO, TNFR1 TKO, and TNFR1 TKO-ADAR1 p150 OE cells were treated with media alone or IFNγ (10 ng/mL) and analyzed by IncuCyte. Values are triplicate mean ± SD. ****P<0.0001 by unpaired t test comparing last data point.

### Extended Data Figure 5

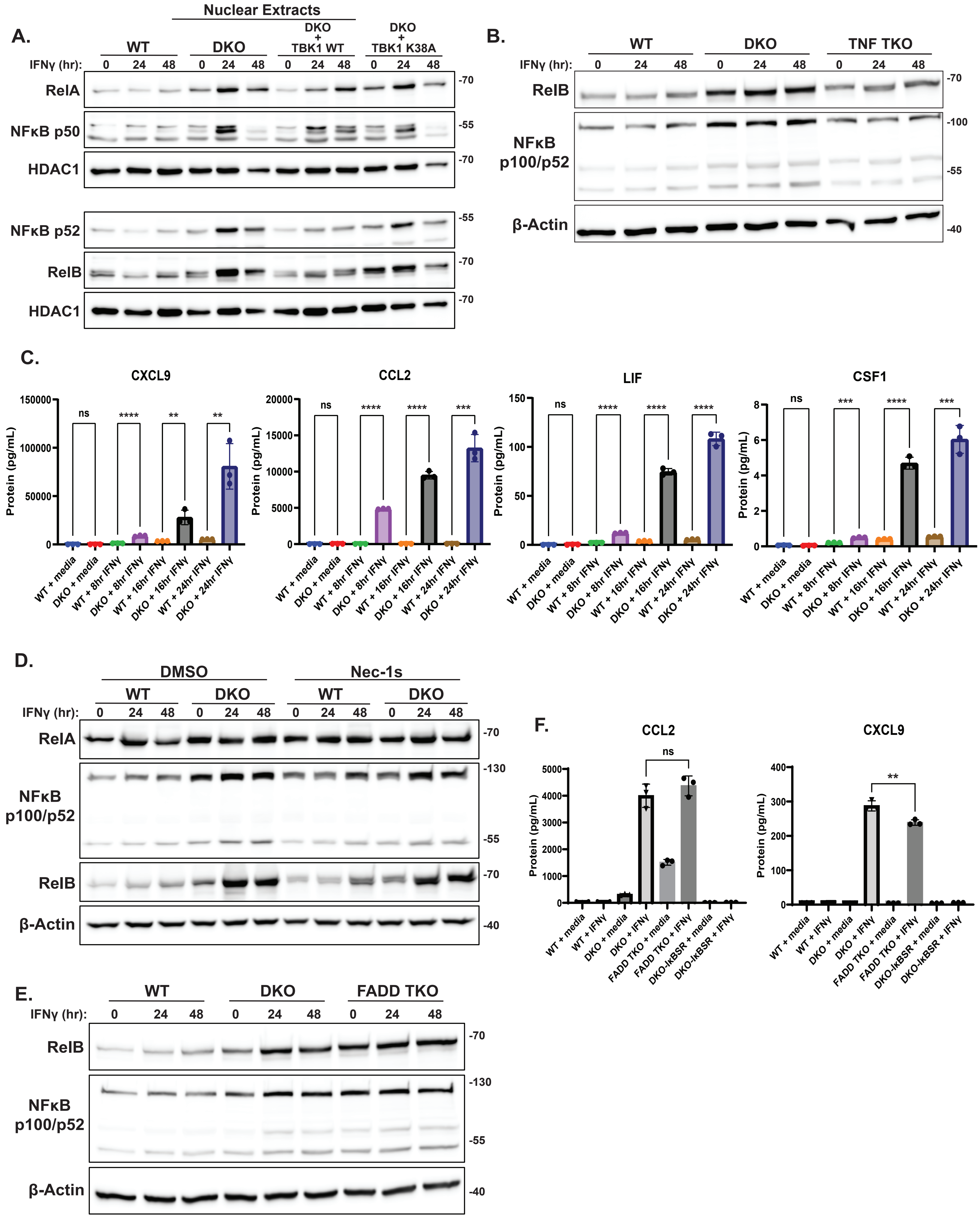

**Extended Data Figure 5. Supporting data for the role of TBK1/IKKε in suppressing NFκB-dependent inflammation in IFNγ-stimulated cells. (A)** Western blot analysis of nuclear extracts in WT, DKO, DKO + TBK1 WT, and DKO + TBK1 K38A B16 cells treated with IFNγ (100 ng/mL) for 0, 24, and 48 h. **(B)** Western blot analysis of the indicated proteins in WT, DKO, and TBK1/IKKε/TNF TKO B16 cells treated with IFNγ for 0, 24, 48 h. **(C)** Luminex analysis of the indicated chemokines in supernatants from WT and DKO B16 cells treated with IFNγ (100 ng/mL) for 0, 8, 16, and 24 h. Statistical analysis was performed using unpaired t test. ns = not significant, **P<0.01, ***P<0.001, ****P<0.0001. **(D)** Western blot analysis of the indicated proteins in WT and DKO B16 cells pretreated with DMSO or Nec-1s (10 μM) for 30 min followed by IFNγ for 0, 24, and 48 h. **(E)** Western blot analysis of the indicated proteins in WT, DKO, and TBK1/IKKε/FADD TKO B16 cells treated with IFNγ for 0, 24, and 48 h. **(F)** ELISA of mouse CCL2 and CXCL9 in WT, DKO, TBK1/IKKε/FADD TKO, and DKO-IκBSR B16 cells treated with IFNγ for 0 and 24 h. ns = not significant, **P<0.01, statistical analysis was performed using unpaired t test.
